# Permissive Fatty Acid Incorporation in Host Environments Promotes Staphylococcal Adaptation to FASII Antibiotics

**DOI:** 10.1101/635698

**Authors:** Gérald Kénanian, Claire Morvan, Antonin Weckel, Amit Pathania, Jamila Anba-Mondoloni, David Halpern, Audrey Solgadi, Laetitia Dupont, Céline Henry, Claire Poyart, Agnès Fouet, Gilles Lamberet, Karine Gloux, Alexandra Gruss

**Affiliations:** Micalis Institute UMR1319, INRA, AgroParisTech, Université Paris-Saclay, Jouy en Josas, France; INSERM U1016, Institut Cochin; CNRS UMR 8104; Université Paris Descartes, Paris, France; SAMM, UMS IPSIT, Faculté de Pharmacie, Université Paris-Saclay, Chatenay-Malabry, France; PAPSSO platform, Micalis Institute, INRA, AgroParisTech, Université Paris-Saclay, Jouy-en-Josas, France; Centre National de Référence des Streptocoques, Hôpitaux Universitaires Paris Centre Site Cochin, APHP, Paris, France

**Keywords:** Conditional antibiotic adaptation, antibiotic resistance, membrane phospholipids, fatty acid stress, triclosan, AFN-1252

## Abstract

Development of fatty acid synthesis pathway (FASII) inhibitors against the major human pathogen *Staphylococcus aureus* hinges on the accepted but unproven postulate that an endogenously synthesized branched chain fatty acid is required to complete membrane phospholipids. Evidence for anti-FASII efficacy in animal models supported this view. However, restricted test conditions used previously to show FASII antibiotic efficacy led us to investigate these questions in a broader, host-relevant context. We report that *S. aureus* rapidly adapts to FASII antibiotics without FASII mutations when exposed to host environments. Treatment with a lead FASII antibiotic upon signs of infection, rather than just after inoculation as commonly practiced, failed to eliminate *S. aureus* from infected organs in a septicemia model. *In vitro*, addition of serum facilitated rapid *S. aureus* FASII bypass by environmental fatty acid (eFA) replacement in phospholipids. Serum lowers membrane stress, leading to increased retention of the two substrates required for exogenous fatty acid (eFA) utilization. In these conditions, eFA occupy both phospholipid positions 1 and 2, regardless of anti-FASII selection. This study revises conclusions on *S. aureus* fatty acid requirements by disproving the postulate of fatty acid stringency, and reveals an Achilles’ heel for using FASII antibiotics to treat infection in monotherapy.

**Significance statement:** Antibiotic discovery to overcome treatment failure has huge socio-medical and economic stakes. The fatty acid synthesis (FASII) pathway is considered an ideal druggable target against the human pathogen Staphylococcus aureus, based on evidence of anti-FASII efficacy in infection models, and the postulate that S. aureus synthesizes an irreplaceable fatty acid. We report that S. aureus alters its behavior in host-relevant conditions. Administering FASII antibiotics upon signs of infection, rather than just after inoculation as frequently practiced, failed to clear septicemic infections. In serum, S. aureus rapidly overcomes FASII antibiotics by incorporating alternative fatty acids. We conclude that previously, premature antibiotic treatments and experimental constraints masked S. aureus antibiotic adaptation capacity. These findings should help streamline future drug development programs.

## Introduction

Fatty acid synthesis (FASII) pathway enzymes are priority targets for ongoing drug development against methicillin resistant *Staphylococcus aureus* (MRSA) (1–11). However, anti-FASII efficacy remains a critical point of debate (12–14). Remarkably, *S. aureus* FASII sensitivity *versus* tolerance hinges on a single issue; whether environmental fatty acids (eFA) can occupy the presumably stringent 2-position of membrane phospholipids (the 1-position is permissive) (7, 14). Anti-FASII resistance due to mutations affecting the target enzyme or to horizontal transfer of an antibiotic-resistant gene homologue may occur and are often antibiotic-specific (15, 16). In contrast, resistant mutants that allow compensatory fatty acid utilization at both phospholipid positions were isolated and also found in clinical isolates; these mutants map in FASII initiation genes distinct from the gene encoding the antibiotic target protein (17–19). Despite emergence of mutations, continued FASII-targeted drug development is rationalized by the accepted postulate that the general wild type *S. aureus* population must synthesize branched chain fatty acid *anteiso* 15:0 (*ai*15) to complete membrane phospholipids (1, 3, 5, 8-11, 14, 20). We investigated this postulate, and report here that an alternative antibiotic adaptation mechanism is functional in host environments, which enables *S. aureus* growth in FASII antibiotics.

## Results and Discussion

### A FASII antibiotic does not clear *S. aureus* infection in a septicemia murine model

Results of animal tests are decisional checkpoints for antibiotic development. FASII antibiotic challenge tests to date administer treatments within minutes to a few hours post-infection (summarized in (19)), *i.e.*, prior to bacterial dissemination to host organs, and before clinical symptoms would call for antibiotic treatment (21, 22). This consideration guided the design of the infection and treatment protocol used here (Fig. 1A). *S. aureus* methicillin resistant strain USA300 was administered by the intravenous route. Antibiotic treatments were initiated 16 h (T16) post-infection, at which time animals exhibited signs of sickness (lethargy and ruffled fur). Group 1 received no treatment. At T16, Group 2 received AFN-1252 a pipeline FASII antibiotic targeting FabI, an enoyl-acyl-carrier-protein-reductase, following recommended dosing (23). Group 3 received vancomycin, which was used to validate that treatment starting at T16 was feasible. At T40 (*i.e.*, 24 h post antibiotic treatment), bacterial counts were significantly lower in organs of both antibiotic-treated animals compared to the untreated group, attesting to AFN-1252 activity (Fig. 1B). In contrast, at T88, whereas vancomycin-treated animals were essentially free of bacteria, all organs from AFN-1252-treated mice still contained *S. aureus* CFUs. Bacterial counts were increased 10-fold in kidneys (to 5×10^6^; p= ≤0.01), decreased 10-fold in liver (p= ≤0.05), and unchanged in spleen. FASII antibiotic treatments thus failed to eliminate *S. aureus* in a septicemia model. The underlying mechanisms leading to FASII inhibitor escape in host-relevant conditions were investigated.

**Fig. 1.**
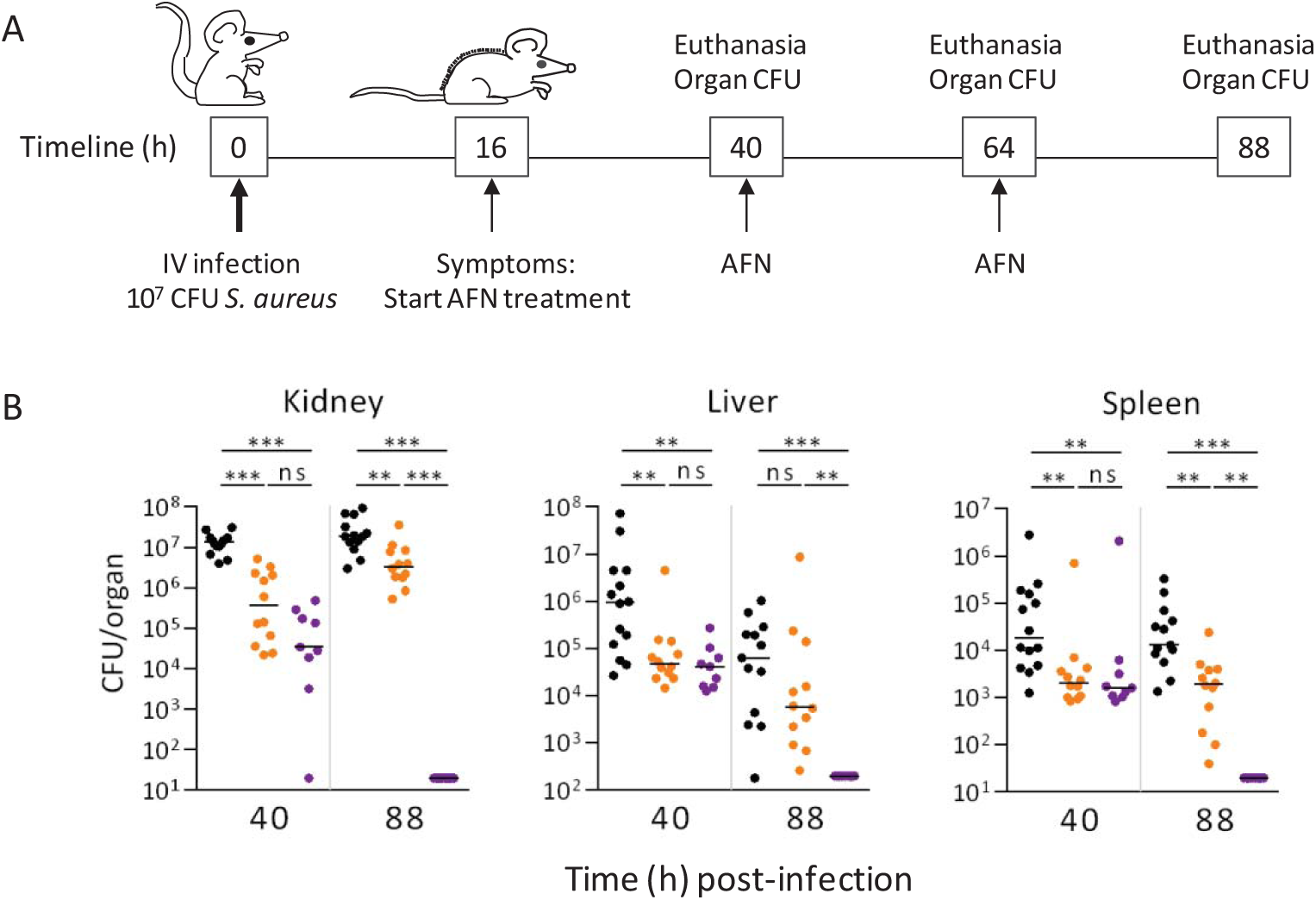
FASII antibiotic efficacy in an *S. aureus* septicemia murine model. **A.** Experimental protocol. Balb/C mice were infected intravenously at T0 with 1×10^7^ CFU USA300. AFN-1252 or vancomycin (control for antibiotic efficacy) was administered at T16 h when mice first showed signs of infection (lethargy, ruffled fur), and at regular intervals thereafter until animals were euthanized. **B.** CFUs were determined in organs upon sacrifice, at T40 h, and T88 h. The above data are pooled from two independently performed experiments. Black lines indicate median CFU corrected to per gram/organ. ***, p<0.001; **, p<0.01; ns, non-significant. Y axis, CFU. Detection limit was 20 CFUs for kidney and spleen, and 200 for liver. Mouse groups are: black, untreated control; orange, AFN-1252-treated; purple, vancomycin-treated.

### Host constituents promote rapid staphylococcal adaptation to FASII antibiotics

We reasoned that in septicemic infection, serum and other host constituents bind eFA, and may neutralize FASII inhibitors (2, 5, 24, 25). The effect of serum on FASII antibiotic activity was tested using triclosan, an extensively studied and used biocide that also targets FabI (26). *S. aureus* triclosan sensitivity was compared in medium containing a 3-fatty-acid cocktail (called here ‘FA’; with triclosan, FA-Tric), and the same medium supplemented with serum (SerFA-Tric)(Fig. 2A). USA300 and Newman strain growth without serum were inhibited by triclosan, with emergence of FASII mutants usually after 24-48 h incubation (18). However, serum supplementation markedly shortened latency compared to BHI cultures, to 8 h and 10 h for USA300 and Newman respectively, and was followed by near-normal growth (Fig. 2A, Fig. S1A). If *S. aureus* outgrowth were due to triclosan titration by serum, FASII would remain active, so that bacterial fatty acid composition would be endogenous. However, the contrary occurred: bacterial fatty acid profiles during outgrowth in SerFA-Tric medium were totally exogenous (Fig 2B, Fig. S1B). Addition of albumin, a major serum constituent, rather than full serum also resulted in FASII bypass (available upon request). Similarly, when USA300 was grown with liver or kidney extracts (without added fatty acids) and triclosan, outgrowth kinetics were similar to those of SerFA-Tric cultures, and cells bypassed the FASII block by incorporating fatty acids from organ lipids (Fig. S2).

**Fig. 2.**
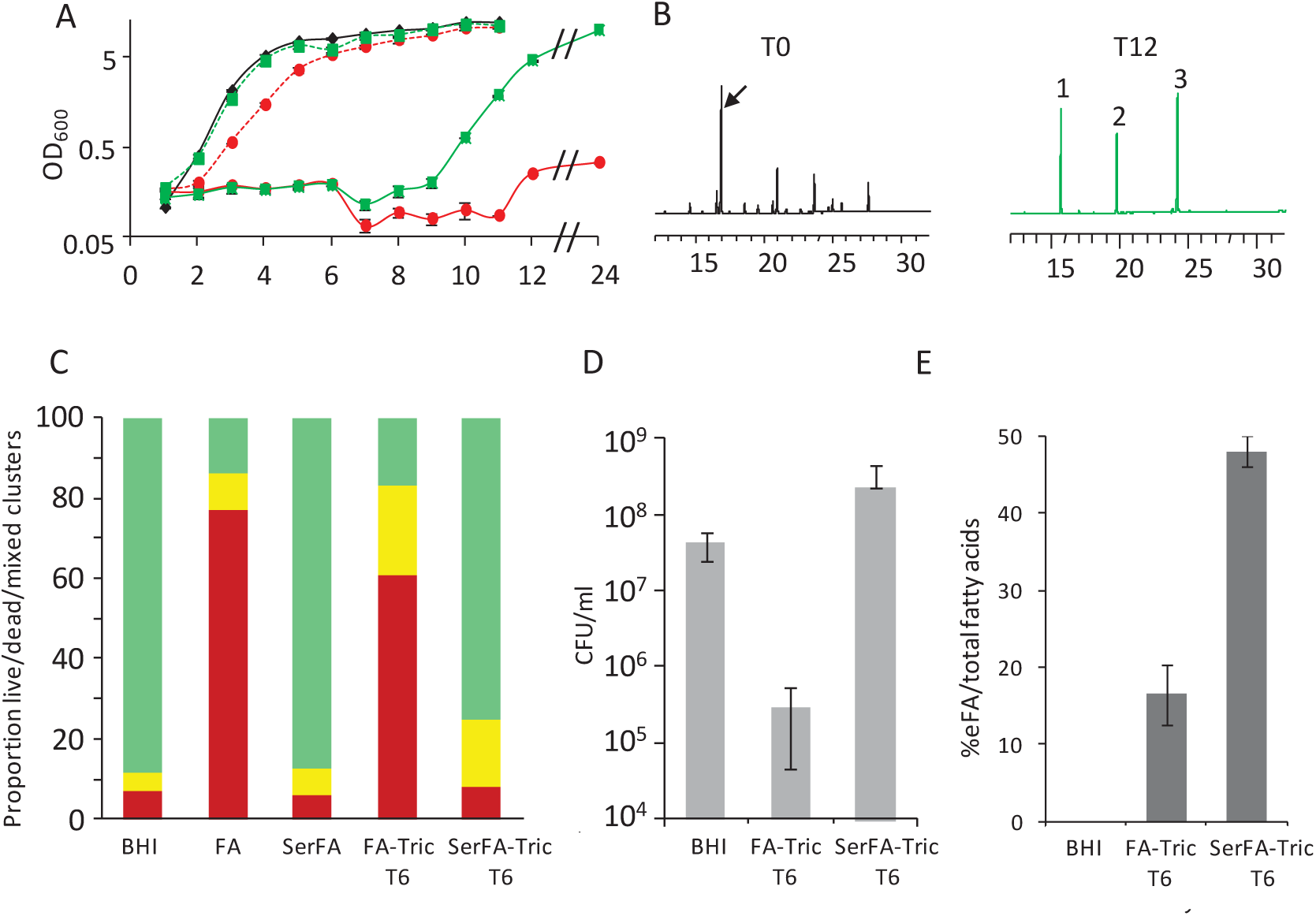
Positive effects of serum on *S. aureus* adaptation to a FASII antibiotic. *S. aureus* strain USA300 was cultured in BHI, FA (a 3-fatty acid cocktail), or SerFA (serum-supplemented FA) media containing or not triclosan (FA-Tric and SerFA-Tric, respectively). **A**. Bacterial optical density (OD_600_). Growth in SerFA-Tric resumed after an ∼8 h latency period, whereas no growth was observed over 24 h in the absence of serum (FA-Tric). Black line, BHI; red lines FA (dashed), and FA-Tric (solid); green lines, SerFA (dashed), and SerFA-Tric (solid). Curves are the average of three biological replicates. **B**. Fatty acid profiles. USA300 in BHI (left), displays endogenous fatty acids; SerFA-Tric–grown cells comprise exclusively eFAs. Black arrow indicates endogenously produced *ai*15. eFA: 1, C14:0; 2, C16:0, 3, C18:1. Representative fatty acid profiles from over 10 determinations are shown. **C.** Cell vitality and permeability were evaluated by fluorescence microscopy using respectively Syto9^TM^ and propidium iodide (PI) probes. USA300 was grown in FA-Tric and SerFA-Tric for 6 h; mid-exponential phase cultures in BHI, FA, and SerFA were used as references. Proportions of vital and permeable bacteria were determined on ∼10,000 bacteria per condition, issued from 3 independent experiments giving comparable images (Fig. S5). Proportions of vital (green), permeable (red), and mixed cell clusters (yellow) are shown. **D.** CFUs were determined in parallel on three independent cultures per condition: BHI starting culture [T0], FA-Tric, and SerFA-Tric samples [T6 = 6 h]). **E.** Fatty acid profiles were determined on 4 biological replicates grown as in ‘C’. Proportions of eFA compared to total fatty acids are shown after 6 h selection. Note that fatty acid profiles are fully exogenous upon outgrowth.

Importantly, pre-incubation in SerFA prior to FASII antibiotic treatments shortened the time prior to *S. aureus* outgrowth. USA300 was challenged with the AFN-1252, which led to a longer (10 h) latency phase prior to outgrowth than did triclosan. However, pre-incubation in serum shortened latency compared to that in non-selective medium to around 6½ h for both drugs (Fig. S3), indicating that bacterial pre-exposure to the lipid-rich host environment contributes to limiting FASII antibiotic efficacy.

*Staphylococcus epidermidis, haemolyticus*, and *lugdugensis* are emerging pathogens that, like *S. aureus*, synthesize branched chain fatty acids (Fig. S4). Representative strains were grown in SerFA and treated with AFN-1252 as above. All cultures grew after overnight incubation and displayed exogenous fatty acid profiles, indicating that these staphylococcal species also bypass FASII inhibitors.

These results show that in serum, *S. aureus* and other staphylococcal species escape anti-FASII inhibition and maintain robust growth by replacing endogenously synthesized fatty acids with eFA. They indicate that serum does not prevent, but actually enhances eFA incorporation by *S. aureus*.

### Adaptation to FASII inhibitors is not due to FASII mutations

Mutations in FASII initiation genes may lead to antibiotic resistance (14, 17-19). We monitored mutations in FASII antibiotic-adapted USA300 or Newman strains by DNAseq, using FASII inhibitors triclosan or AFN-1252 (Table S1). DNAseq of USA300 grown in BHI and SerFA, and Newman grown in BHI were used as references. Adaptation was confirmed by exogenous fatty acid profiles of antibiotic-grown samples in 12-14 h cultures. Eight of nine adapted strains carried wild type FASII genes (the exception was mutated in SAUSA300_1476 encoding FASII initiation gene *accB*). One isolate displayed no detectable genome mutations. The other clones carried SNPs corresponding to commonly found variants and are thus likely unrelated to FASII antibiotic adaptation (described in Table S1). The absence of FASII mutations in FASII-antibiotic-adapted clones in serum distinguishes this adaptation mechanism from resistance due to FASII mutations (15, 18). *S. aureus* evasion of FASII antibiotics in serum identifies a novel strategy of condition-dependent adaptation.

### Serum lowers fatty-acid-induced bacterial membrane permeability and improves fitness

Numerous fatty acids perturb bacterial membrane integrity and are a source of stress, while serum albumin neutralizes these effects (24, 25, 27, 28). Accordingly, serum abolished the eFA-provoked growth lag in non-selective medium (Fig. 2A). Serum effects on *S. aureus* cell vitality and permeability, and cell state, were examined. Free fatty acids had strong permeabilizing effects on cells from FA and FA-Tric, as compared to BHI cultures, as evaluated by fluorescence microscopy; these effects were offset by serum in SerFA and SerFA-Tric cultures (Fig. 2C, Fig. S5A, S5B). Plating efficiency was ∼10^3^-fold higher after 6 h growth in SerFA-Tric compared to FA-Tric (Fig. 2D). The accumulation of tetrads comprising mixed-stained cells in triclosan-treated cultures suggested a cell division block, which could explain the observed latency prior to outgrowth (Fig. S5C). Importantly, serum facilitates fatty acid incorporation in the latency period, as seen in 6 h FA-Tric and SerFA-Tric cultures (Fig. 2E). The bacterial stress state in FA-Tric supplemented or not with serum was also assessed by a proteomics approach (Fig. S6). Differences in stress-related protein abundance between anti-FASII-treated *versus* control cultures were overall more pronounced when serum was absent. Serum therefore improves *S. aureus* fitness in fatty acid-containing environments and contributes to FASII antibiotic adaptation *via* eFA incorporation.

### Retention of the FASII bypass precursors acyl carrier protein (ACP) and eFA is increased in serum

We questioned how serum affected availability of two key substrates required for FASII antibiotic adaptation, ACP and eFA. ACP is required for both *de novo* fatty acid synthesis *via* FASII, and for eFA incorporation in the phospholipid 2-position (Fig. 3A) (18, 29). eFA induce membrane leakage that depletes *S. aureus* ACP pools, and could limit eFA incorporation during FASII inhibition (14, 28). *S. aureus* ACP pools were compared by immunoblotting using anti-ACP antibodies in total extracts from cultures grown without and with serum and triclosan (Fig. 3B). ACP levels were lower in extracts from cells grown in eFA compared to BHI medium, as reported (28). ACP was barely detected in FA-Tric-grown cells after 2 h, and remained low at 4 h. In contrast, serum addition reversed this effect, leading to greater intracellular ACP availability.

**Fig. 3.**
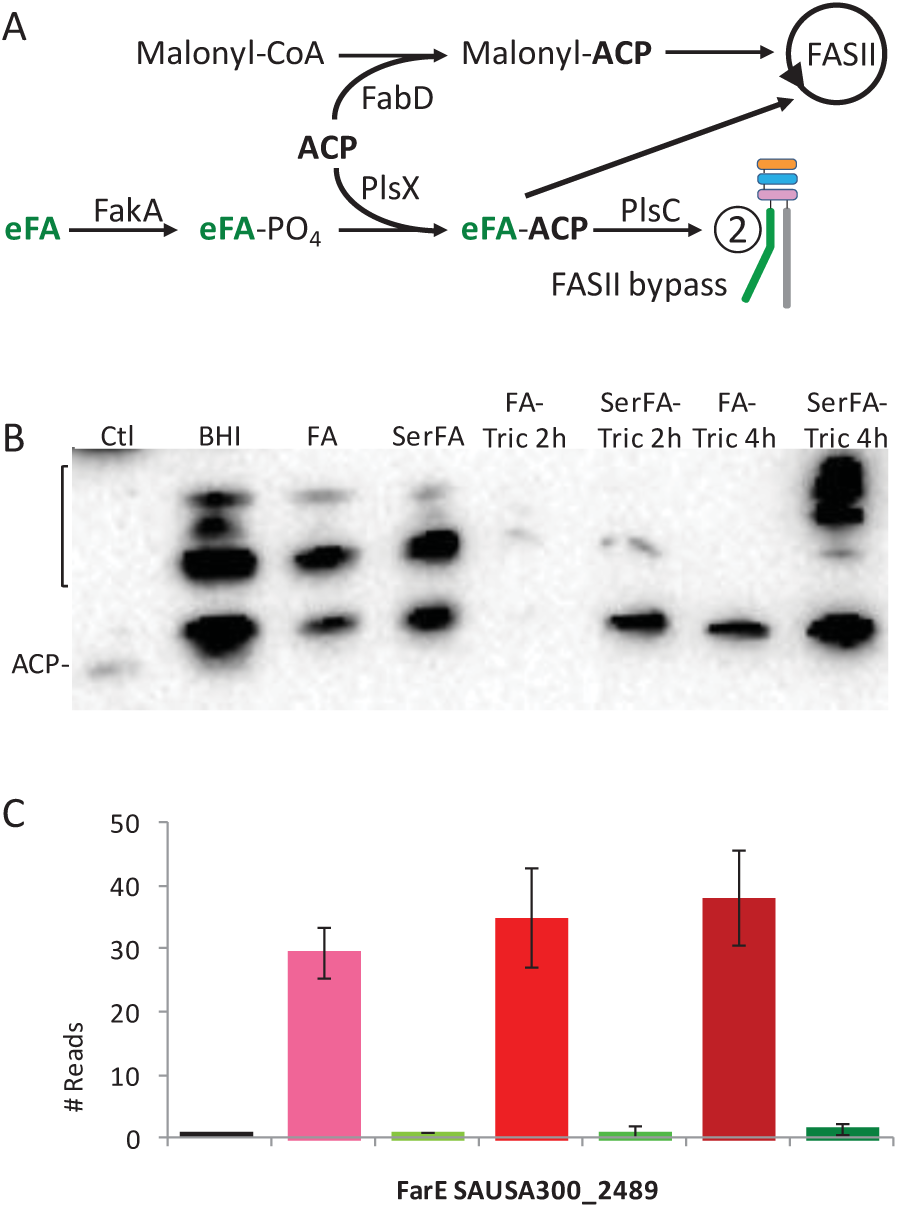
Impact of serum on *S. aureus* intracellular retention of ACP and eFA, and eFA replacement in phospholipids. **A.** ACP and eFA (green) are required substrates for FASII bypass, *via* eFA incorporation in the phospholipid 2-position (lower right). FASII and FASII bypass enzymes compete for ACP and eFA-ACP intermediates. **B.** ACP immunodetection. Cell extracts were prepared from *S. aureus* USA300 *spa* strain. BHI, FA, and SerFA cultures were harvested at OD_600_ = 1, and triclosan-treated cultures were harvested at 2 h and 4 h. Equivalent protein concentrations were loaded on SDS-PAGE gels. Immunoblotting was performed using anti-ACP antibody. Migration of purified ACP is shown at left. Multiple bands (bracket) may correspond to non-dissociable acyl-ACP-protein complexes. **C.** Expression of fatty acid efflux pump FarE. Proteomic analyses were performed on *S. aureus* USA300 *spa* treated as in ‘B’. Samples are in the same order as in ‘B’. Black, BHI; pink, FA; light green, SerFA; red, FA-Tric 2h; middle-green, SerFA 2h; burgundy, FA-Tric 4h; dark green, SerFA 4h. Quadruplicate independent samples were used for immunoblots and proteomic analyses. The full proteomic study is available at doi:10.17632/9292c75797.1.

Intracellular eFA pools are limited by a recently characterized *S. aureus* fatty acid efflux pump, FarE, which is induced by unsaturated eFA (30, 31). Proteomics analysis (described above) confirmed high FarE expression in all eFA-containing media without serum. In contrast, nearly no FarE was detected in cells issued from serum-containing cultures despite the presence of eFA (Fig. 3C). Greater *S. aureus* intracellular retention of ACP and eFA in serum is consistent with more efficient eFA incorporation and FASII antibiotic adaptation (Fig. 2E).

### Exogenous fatty acids are incorporated in the phospholipid 2-position in the absence and presence of FASII antibiotics

The accepted but unproven rationale for developing *S. aureus* FASII inhibitors is the stringent requirement for endogenous branched chain fatty acid *ai*15 at the phospholipid 2-position, catalyzed by the PlsC acyltransferase (1, 2, 10, 14, 32). The above results gave evidence against the generality of phospholipid stringency by showing that FASII antibiotic adaptation leads to eFA incorporation at both phospholipid positions (Fig 2B, Fig. S1B). However, eFA incorporation in both *S. aureus* phospholipid positions could be a last resort choice when the preferred substrate *ai*15 is unavailable. Alternatively, the use of eFA *versus ai*15 may simply depend on intracellular substrate availability, which increases in serum. eFA incorporation was assessed in non-selective conditions to discriminate between these alternative hypotheses. To remove ambiguity in distinguishing endogenous from exogenous fatty acids in phospholipid identifications, a cocktail of unsaturated eFA 17:1*trans* (*tr*) and 18:1*cis* prepared in delipidated serum (dSer2FA medium) was used to supplement *S. aureus* USA300 growth. Although structurally distinct from *S. aureus* endogenous fatty acids, 17:1*tr* and 18:1*cis* did not interfere with growth in the serum-containing medium (OD_600_ = 6.4 for both BHI and dSer2FA cultures at 6 h). In this non-selective growth condition, 17:1*tr* and 18:1*cis* comprised about 60% of the total fatty acid content (Fig. 4A, Fig. S7A) at the expense of straight chain saturated fatty acids, which decreased from 50% in BHI, to about 10% in dSer2FA cultures. Importantly, eFA occupied both positions in phosphatidylglycerol (PGly) phospholipids after 6 h without antibiotic (Fig. 4B, Table S2). These results prove that wild type *S. aureus* incorporates dissimilar fatty acids in both phospholipid positions without the need for FASII antibiotic selection, and rule out previously assumed fatty acid selectivity of phospholipid-synthesizing enzymes.

**Fig. 4.**
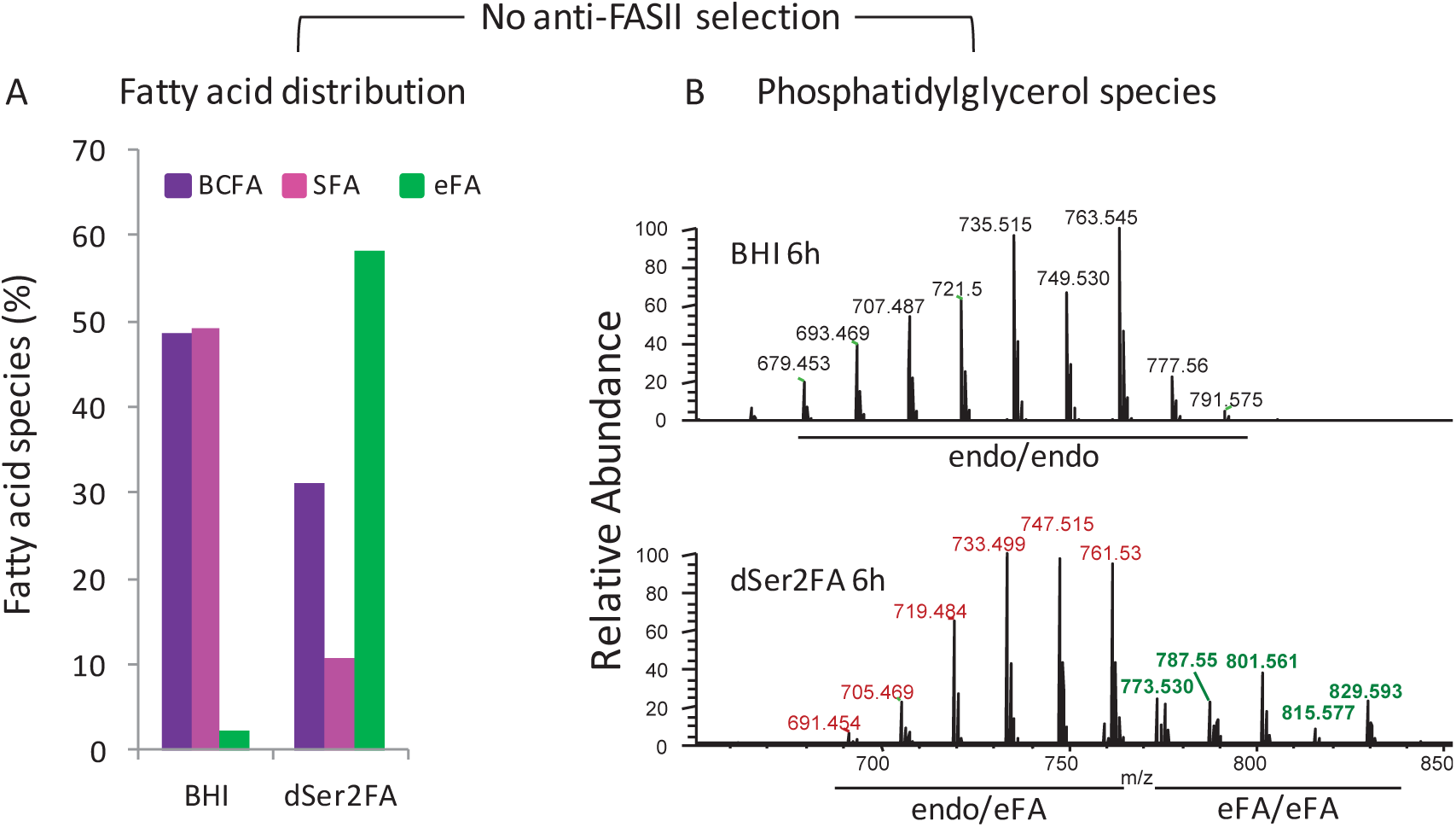
Serum promotes eFA replacement in phospholipids without antibiotic selection. *S. aureus* USA300 was grown 6 h in 10% delipidated serum containing equimolar C17:1*tr* - C18:1*cis* mix (dSer2FA). **A.** FA composition. Fatty acid species are presented as the proportion of endogenous branched chain fatty acids (BCFAs C15:0, i15, *ai*15; purple), endogenous saturated (straight chain) fatty acids (SFAs C18:0, C20:0; pink), and eFA (C17:1*tr* and C18:1*cis*, green); see Fig. S7A for fatty acid profiles. **B.** Phosphatidylglycerol (PGly) MS profiles from samples in ‘A’: in black, masses of PGly species with endogenous fatty acids in both positions (endo/endo); in red, PGly species with one endogenous fatty acid and one eFA (endo/eFA); in green, PGly species with eFA in both positions (eFA/eFA). Without antibiotic, 20-25% of PGly species comprise eFA at both positions as estimated from peak heights. See Table S2 for fatty acids comprising major PGly species.

Fatty acid and PGly profiles were determined 4, 6, 8, 10, and 15 h after USA300 treatment with AFN-1252 or triclosan, in parallel with OD_600_ (Fig. S7B, Fig. S8). For both FASII antibiotics, bacterial transition from mixed to exclusively exogenous fatty acids was concomitant with outgrowth from latency starting at 8 h post-treatment, and was completed at 10 h (Figs. S8A, S8C). For both FASII antibiotic treatments, predominant phospholipid species at 10 h were totally exogenous (18:1*cis*|18:1*cis*, 17:1*tr*|17:1*tr*, and 18:1*cis*|17:1*tr*; Fig. S8B and S8D, Table S3). The sharp increase in exclusively eFA-containing phospholipids coincides with the exit from latency, as expected from coordination between membrane phospholipid synthesis and cell growth (33, 34). These results show that when *S. aureus* membrane integrity is maintained, as by host constituents, eFA incorporation is not stringent, and reflects competition between fatty acids synthesized by *S. aureus* and those available from the environment. eFA are incorporated in both phospholipid positions in the absence of selection, and completely replace endogenous fatty acids in the presence of FASII antibiotics.

## Conclusions

*S. aureus* is shown here to to adapt to FASII antibiotics in host environments and without FASII mutations. A FASII antibiotic was ineffective in a septicemia model in which the treatment protocol respected the interval between primo-infection and treatment time. Design of antibiotic challenge tests based on realistic intervals between infection and treatment should improve the predictive value of animal studies, which are decisive for scale-up to clinical trials. Based on this study, AFN-1252 and likely any FASII inhibitor are likely ineffective as stand-alone treatment of *S. aureus* deep infection.

*S. aureus* responses to FASII antibiotics are summarized in a model, in which greater bacterial membrane integrity and intracellular ACP and eFA pools in the presence of host components such as serum, lead to complete eFA replacement in phospholipids (Fig. 5). FASII antibiotic adaptation occurs in proliferating bacteria. The rapid kinetics, absence of FASII mutations during antibiotic adaptation, and eFA-replete phospholipids in the absence of selection, indicate that this event impacts a general *S. aureus* population. Serum and other host components promote low stress and FASII adaptation, which links this model to anti-FASII treatment failure. In contrast, skin surfaces producing fatty acids are linked to high stress environments unfavorable to FASII bypass except by FASII mutation (Fig. 4; (8, 18)). FASII-targeting has been recently suggested for *Clostridium difficile* and *Listeria monocytogenes* (35, 36). Like *S. aureus*, these pathogens might show relaxed phospholipid stringency in host environments; niche-dependent adaptation in all these cases remains to be tested.

**Fig. 5.**
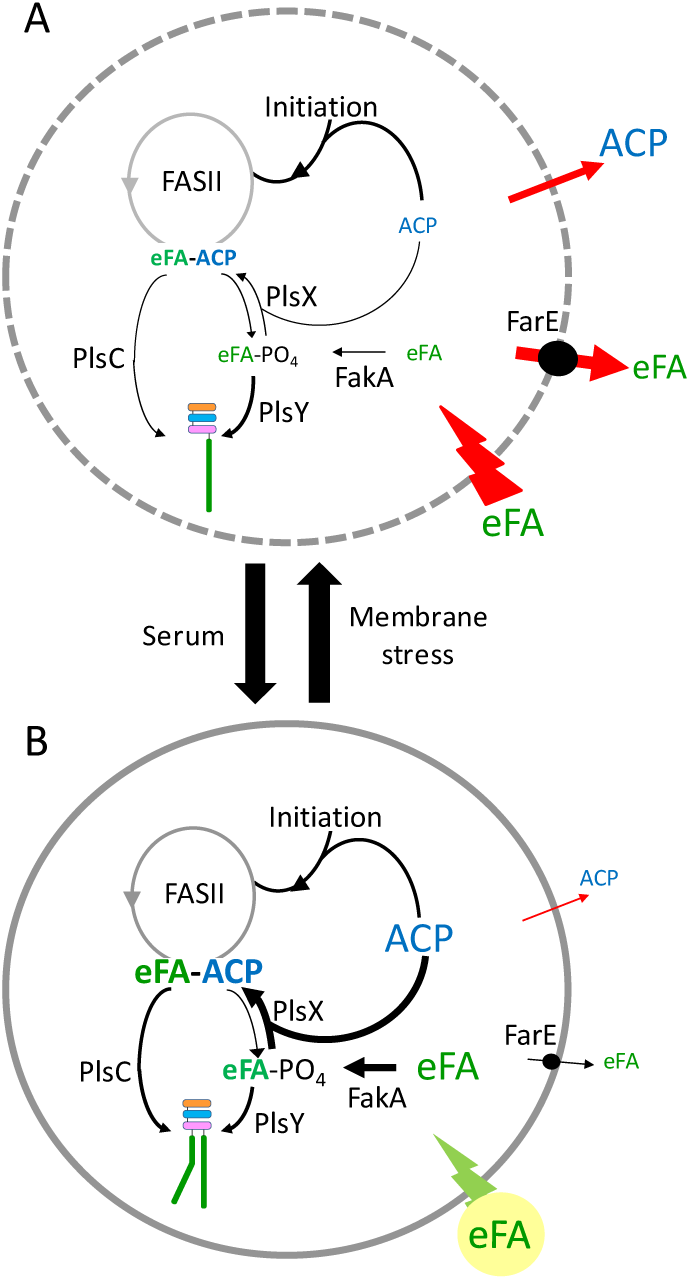
Conditional *S. aureus* adaptation to FASII antibiotics. The membrane stress state impacts eFA and ACP intracellular pools and dictates FASII antibiotic adaptation as shown in the model. **A.** High stress: Free eFA permeabilize membranes. Fatty acid efflux *via* FarE and ACP leakage (28, 30, 31) lead to depletion of FASII-bypass substrates. In this condition, eFA-PO_4_ is used by PlsY to charge the phospholipid 1-position. FASII antibiotics would arrest growth, leading to cell death or emergence of FASII initiation mutants that bypass FASII (17, 18). **B.** Low stress: Serum (yellow disk) or other host components reduce eFA toxicity ((24), this report), and improve membrane integrity. Higher ACP and/or eFA-PO_4_ pools drive PlsX directionality to eFA-ACP production. eFA-ACP, and endogenous acyl-ACP (if not blocked by anti-FASII), compete for phospholipid synthesis at the 2-position *via* PlsC. eFA occupy both phospholipid positions even without FASII antibiotics (Fig. 4B). eFA (green) and ACP (blue) abundance is represented by font size; phospholipids, “π” form; thick arrows, favored reactions; thin arrows, reduced reactions. Dashed circle, permeable membrane; solid circle, intact membrane. PlsY mediates eFA incorporation in the phospholipid 1-position; PlsX and PlsC, catalyze fatty acid insertion in the phospholipid 2-position. Only eFA processing is presented.

Bacterial metabolism and environmental stress play unquestionable roles in the outcome of antimicrobial treatments. For example, reduced metabolism in bacterial persisters is recognized as an major means of escape from antibiotic killing (37, 38). In contrast FASII antibiotic escape involves a latency phase followed by adaptation and robust outgrowth *via* compensatory incorporation of host fatty acids. Such metabolic rescue is likely frequent, as bacteria commonly salvage metabolites from their environments. This consideration may set a logical limit when selecting targets for antimicrobial drug development.

As shown here, *S. aureus* and other Firmicute pathogens may have reduced genetic requirements in host biotopes, including nonessentiality of FASII. Interestingly, a wall-less L-form *S. aureus* multiplies in the presence of cell wall antibiotics and survives by oversynthesis of fatty acids ((39) and references therein). An intriguing possibility is that such “primitive bacteria” can use the lipid supply of the host, taking their lifestyle one step further towards antibiotic adaptation and parasitism.

## Acknowledgements

Nebraska Transposon Mutant Library strains were generously provided by the Network on Antimicrobial Resistance in *Staphylococcus aureus*, prepared by BEI Resources, NIAID, NIH, USA. We are grateful to P. Casedesus (Univ. Sevilla, Spain), B. Michel and P. Bouloc (I2BC, France), and P. Trieu-Cuot (Institut Pasteur, France) for stimulating discussion of this work. We thank M. Gaillard, A. Picart (I. Cochin), and M. Benard (animal facility, I. Cochin) for expert assistance in animal studies, and M. Gaillard for data analyses; M. Gohar (Micalis) for advice in fluorescence microscopy studies; R. Boudjemaa, E. Borezée-Durant, R. Briandet, J. Deschamps, D. Lechardeur, and P. Gaudu (Micalis) for frequent discussion. GK was funded by the French ministry; AP received a postdoctoral grant from the regional DIM-Malinf; LD, AW, and M. Gaillard were funded by the Agence Nationale de la Recherche (StaphEscape project ANR-13001038). We thank the ANR and the Fondation pour la Recherche Medicale (DBF20161136769) for project funding.

## Author Contributions

Physiology, molecular biology, microscopy, and immunodetection tests; GK, CM, AP, JA-M, DH, LD, KG, and AG. Lipid analyses: GL, AS, KG. Proteomics analyses: GK, JA-M, and CH. Animal experiments: AW, AF, CP. Data analyses: GK, CM, AW, AP, JA-M, CP, AF, GL, KG, and AG. Experimental design and project conception: AW, AF, CP, GL, KG, and AG. AG directed the project and wrote the manuscript.

## Declaration of Interests

The authors declare no competing interests.

## Supplementary information

### Materials and Methods

#### Bacterial strains

*S. aureus* RN4220 and Newman strains are from our laboratory collection. *Staphylococcus epidermidis* ATCC 12228, *Staphylococcus haemolyticus* JCSC1435, and *Staphylococcus lugdunensis* N920143 were kindly supplied by R. Briandet and E. Borezée from this institute. Methicillin-resistant USA300 FPR3757 (called here USA300), and the Nebraska library of transposon insertions (University of Nebraska Medical Center) was generously supplied by BEI resources (1). Tested USA300 derivatives from the Nebraska library contained insertions in the following genes: *spa* (SAUSA300_0113 encoding immunoglobulin G binding protein A), SAUSA300_0226, SAUSA300_0242, SAUSA300_0407, SAUSA300_1177, and SAUSA300_1684 (see Table S1).

#### Growth media

BHI medium was used for *S. aureus* growth except in media containing liver extracts. Kidney and liver extracts were prepared as described (2) and used at 3% final concentration and used in BHI or LB medium. Fatty acids (Larodan, Sweden) were prepared as 100 mM stocks in dimethyl sulfoxide (DMSO). The three fatty acid mixture (referred to as « FA ») comprising C14:0 (myristic acid), C16:0 (palmitic acid), and C18:1*cis* (oleic acid) was prepared in a 1:1:1 ratio (0.17 mM each). The two fatty acid mixture comprising C17:1*trans* and C18:1*cis* was used at a 1:1 ratio (0.25 mM each); the more rigid 17:1*trans* species was added to limit membrane fluidity due to C18:1*cis*. Newborn calf serum (Sigma-Aldrich, France), or delipidated calf serum (Eurobio, France) was added to growth medium (10% final concentration) as indicated. Triclosan (referred to as “Tric”; Irgasan; Sigma-Aldrich) was added at 250 ng/ml and 500 ng/ml in medium without and with serum respectively; this corresponds to 15-30 times the minimum inhibitory concentrations (MIC) as determined on BHI medium (2). AFN-1252 (referred to as “AFN”; MedChem Express) was added at 500 ng/ml in all media, which corresponds to about 100-fold the reported *S. aureus* MIC (∼4-8 ng/ml; (3, 4)). FASII inhibitors were prepared in DMSO (1 mg/ml) for all *in vitro* experiments.

#### Infection and antibiotic treatments in mouse septicemia model

Animals were housed in the Institut Cochin animal facility accredited by the French Ministry of Agriculture for performing experiments on live rodents. Animal experimentation was performed in compliance with French and European regulations on care and protection of laboratory animals (EC Directive 2010/63, French Law 2013– 118, February 6, 2013). All experiments were approved by the Ethics Committee of the Paris-Descartes University (agreement n° 2015032714098562). Six-week-old Balb/C mice were inoculated intravenously in the ophthalmic plexus with exponential phase USA300 (1×10^7^ CFU: colony forming units) in 200 µl PBS. Mice were randomized into three groups of at least 18 mice per type of treatment per experiment. Group 1 was mock-treated with excipient (see Group 2); n=14, 6, and 13 for 40 h, 64 h, and 88 h dissections respectively. Group 2 mice received anti-FASII AFN-1252 (Clinisciences France), administered at 16 h, and every 24 hours until sacrifice, by gavage, as reported in all published work using 5 mg/kg (>500 times the reported MIC of 4-8 ng/ml; (3, 4)), delivered in 50 µl as a 2 mg/ml emulsion in 20% PEG 3350; n=12, 6, and 12 for 40 h, 64 h, and 88 h dissections respectively. Group 3, was treated with vancomycin (100 mg/kg in NaCl 0.9%), and administered in 100 µl by intraperitoneal injection at 16 h, and every 12 hours until sacrifice, as described (100 times the MIC determined as 1 µg/ml; (5, 6); n=9, 6, and 12 for 40 h, 64 h, and 88 h dissections respectively. This positive control confirmed the capacity to clear infections when treatment is administered 16 h post-infection. Kidney and spleen were dissected and homogenized (‘hard’ setting, 3 times 20 s) in 1 ml saline (Precellys, Bertin Instruments, France); liver was turraxed in 10 ml saline. Sample dilutions were plated on solid BHI medium for CFU determinations.

#### Determination of *S. aureus* fatty acid profiles

Aliquots of *S. aureus* cultures (routinely an OD_600_ equivalent ≥1 was used) were centrifuged and washed once in 0.9% NaCl containing 0.02% Triton X-100, followed by two washes in 0.9% NaCl. Whole cell esterified fatty acid determinations were then done as described (7). Briefly, cell pellets were treated with 0.5 ml of 1N sodium methoxide in methanol. Heptane (200 µl) was then added, together with methyl-10-undecenoate (Sigma-Aldrich) as internal standard, vortexed for 1 min, and centrifuged. Fatty acid methyl esters were recovered in the heptane phase. Analyses were performed in a split-splitless injection mode on an AutoSystem XL Gas Chromatograph (Perkin-Elmer) equipped with a ZB-Wax capillary column (30m × 0.25 mm × 0.25 µm; Phenomenex, France). Data were recorded and analysed by TotalChrom Workstation (Perkin-Elmer). *S. aureus* fatty acid peaks were detected between 12 and 32 min of elution, and identified with retention times of purified esterified fatty acid standards.

#### Fluorescence microscopy

Bacteria were grown in BHI, FA, Ser-FA to OD_600_ = 0.5-1, or FA-Tric and SerFA-Tric for 6 hours. Cells (OD_600_ equivalent of 10-20) were centrifuged and washed once in phosphate buffered saline. Syto9^TM^ (0.5 µl sample of a 5 mM solution in DMSO; Thermo Fischer Scientific) and propidium iodide (3 µl of 1.5 mM solution in water; Sigma, France) were added to 30 µl samples for respectively viable and permeabilized cell visualization. Cells were observed 10 min post-staining by fluorescent microscopy using a Zeiss AxioObserver Z1 inverted fluorescence microscope equipped with a Zeiss AxioCam MRm digital camera and Zeiss fluorescence filters. Images were processed with the Zeiss ZEN software package using a 38 HE Green Fluorescent Protein filter (excitation wavelength 450/490 nm; beam splitter, 495 nm; emission, 500/550 nm) and 45 Texas Red filter (excitation: 540/580 nm; beam splitter, 585 nm; emission, 595-668 nm). The numbers of live (green) and permeabilized (red) cells were counted manually. Dividing cells and tetrads were counted as single entities. Tetrads and clusters containing both viable and permeabilized cells (in FA-Tric and SerFA-Tric samples) were classified in a separate “mixed” category. The proportion of tetrads among total cells was determined by manual counting; 15 micrographs from 3 independent cultures per condition were evaluated, based on a total of ∼10 000 cell clusters per condition.

#### Genome sequencing

*S. aureus* USA300 and Newman strains were harvested after exit from latency phase for cultures grown in SerFA-Tric (12 h cultures; 3 independent samples for each strain), or in SerFA-AFN (15 h cultures; 3 independent samples of USA300). Cultures in BHI (one sample for each strain), and in SerFA (for USA300) were sequenced as references. DNA extractions were performed using the Qiagen “DNeasy_®_ Blood & Tissue Kit”, following manufacturer’s protocol, except that cell pellets were first resuspended in 0.1 mg lysostaphin/ml Tris 10 mM (AMBI, USA) and incubated 30 minutes at 37°C. Genomic DNA sequencing by Illumina HiSeq next generation sequencing was outsourced (GATC-Biotech, Konstanz, Germany). Coverage was estimated to be at least 70-fold for USA300 SerFA-Tric samples, and at least 400-fold for all other samples. The 2 × 150 paired-end reads were analysed using “Variation Analysis” method provided by Patricbrc.org. Bowtie2 (Patricbrc.org) was used to align sequences and SAMtools to identify SNPs (8). SNPs that differed in non-antibiotic-treated USA300 and Newman strains from those in the reference sequence (GenBank Nucleotide accession codes NC_007793.1 and NC_009641 respectively) were subtracted prior to variant screening. Variants were identified as representing at least 80% of reads in sequences for which there were at least 10 reads.

#### ACP assessment by immunoblotting

The USA300 *spa::Tn* strain (SAUSA300_0113; (1)) was used for immunoblotting to avoid IgG titration by Protein A; this strain was confirmed to behave like its parent with respect to FASII antibiotics. An overnight BHI USA300 *spa* preculture was used to inoculate BHI, FA, SerFA, FA-Tric, or SerFA-Tric media at OD_600_ = 0.1. Cultures were harvested at OD_600_ = ∼1 for BHI, FA, and SerFA, and after 2 h or 4 h for FA-Tric or SerFA-Tric. All samples were adjusted to equivalent OD_600_ values, and washed twice in TE-protease inhibitor (cOmplete Tablets, Mini *EASYpack* Roche, Germany, as per supplier’s instructions), prior to lysis with Fastprep. Samples (20 µg per well as quantified by the Bradford Protein Assay kit (BioRad)) were treated for 3 min at 95°C and then loaded on 12.5% SDS-PAGE gels run at 150 volts for 2 h. Gels were then electro-transferred to PVDF membranes (0.2 mm; BioRad; 75mA) for 3 h on a semi-dry transfer unit (Hoefer TE 70). Western blotting and exposure used an ECL kit (Perkin-Elmer) as per supplier’s instructions. Rabbit anti-*S. aureus* ACP antibodies (2) were used at 1:1,300 dilution.

#### Proteomic analyses of USA300 responses to anti-FASII

Four independent overnight BHI precultures of USA300 *spa::Tn* strain were used to inoculate BHI, FA, SerFA, FA-Tric, or SerFA-Tric media at OD600 = 0.1. Culture extracts were prepared as described for Western blotting, and after verification of protein concentration and quality, 10 μg of each protein sample was short-run on SDS-PAGE. Further sample treatment by LC-MS/MS, and bioinformatics and statistical analyses of data are as described (9). The reference genome GenBank Nucleotide accession code NC_007793.1 was used for protein annotation. The complete list of proteins expressed in the five growth conditions is available on the Mendeley database (doi:10.17632/9292c75797.1; https://data.mendeley.com/datasets/9292c75797/draft?a=bb34343b-6314-4421-89bd-b25e2bbdb0df) upon publication.

#### Extraction of polar membrane lipids

Lipid extractions were performed as described with modifications (10, 11). Briefly, freeze-dried cell material (100 mg) was extracted with 9.5 ml of chloroform-methanol 0.3% NaCl (1:2:0.8 v/v/v) at 80°C for 15 min. All following steps were done at room temperature. Extracts were vortexed for 1 h and centrifuged for 15 min at 4000 rpm. Supernatants were collected and cell debris was re-extracted with 9.5 ml of the same mixture, vortexed for 30 min, and centrifuged. Supernatants were then pooled and 5 ml each of chloroform and 0.3% NaCl was added and mixed. Phase separation was achieved by centrifugation at 4000 rpm for 15 min. The upper phase was discarded and the collected chloroform phase was evaporated to dryness under a nitrogen stream and stored at −20°C.

#### Phosphatidylglycerol identification

The polar membrane lipid samples were injected in chloroform in a chromatographic system (ThermoFisher Scientific) including a Dionex U-3000 quaternary RSLC, a WPS-3000RS autosampler and a column oven. Lipid separation was carried out by liquid chromatography on a PVA-Sil column (150 × 2.1 mm I.D., 120 A) (YMC Europe GmbH) with a 10×2 mm guard column packed with the same material. Column temperature was thermostatically controlled at 35°C. The chromatographic method separates phospholipids according to class, and was performed as described (12, 13).

For phosphatidylglycerol species identification, this system was coupled to an LTQ-Orbitrap Velos Pro (ThermoFisher) equipped with an H-ESI II probe. Spray voltage was set at 3.3 kV. Heater temperature of the probe was set at 200°C. Sheath gas, auxiliary gas and sweep gas flow rates were set at 20, 8 and 0 (arbitrary unit) respectively. Capillary temperature was set at 325°C and S-lens RF level at 60%. Analysis was performed in negative mode to obtain structural information on phosphatidylglycerol fatty chains. The mass spectrometer is equipped with two analyzers: a double linear ion trap (LTQ Velos Pro) for fragmentation at low resolution and an orbital trap (Orbitrap©) for high resolution detection. Detection was performed in full MS Scan with 100,000 resolution and data dependent MS^2^ and MS^3^ with collision induced dissociation (CID; collision energy set at 35). Chromatographic retention time was used for polar head identification by comparing to commercial standards. Phospholipid identification was performed using high resolution mass full scan to obtain the formula of the entire PG species (14), and MS^2^/MS^3^ fragmentation to obtain structural information about fatty acid chain composition of each species.

### Statistical analysis

Means and standard errors of replicate growth curves and proportions of fatty acids were determined (Excel, Microsoft, USA). For phospholipid determinations, the average of two experiments with the range of duplicates is presented. Animal experiments data were statistically analyzed by the Mann-Whitney non-parametric test with GraphPad Prism 5.0 (GraphPad Software, San Diego, California).

### Data availability

Genome sequence data have been deposited in the European Nucleotide Archive and can be accessed at http://www.ebi.ac.uk/ena/data/view/PRJEB24433. The complete list of proteins from the proteomic study will be made available on the Mendeley database (doi:10.17632/9292c75797.1) upon publication. The authors declare that all other data supporting the findings of this study are available within the article and its Supplementary Information files, or from the corresponding author upon request.

---

**Fig. S1.**
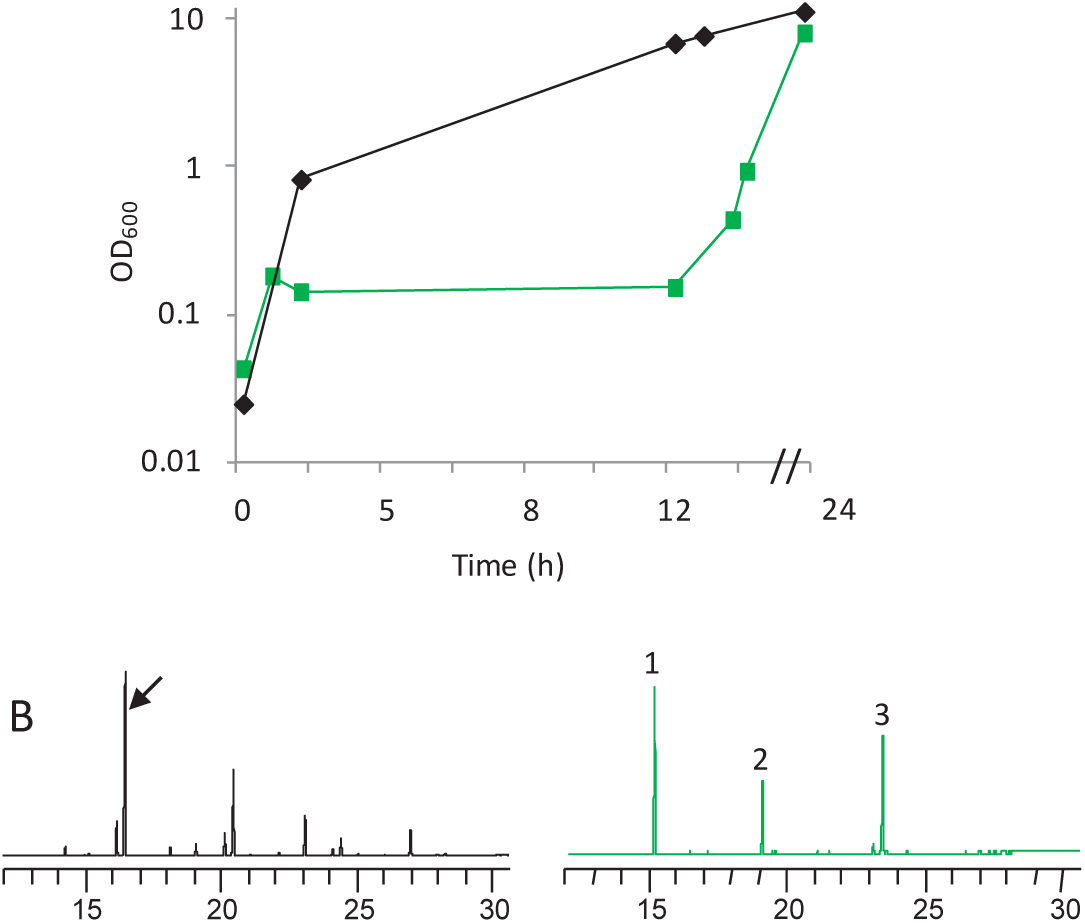
Positive effects of serum on *S. aureus* adaptation to a FASII antibiotic. *S. aureus* Newman strain was cultured in BHI and SerFA-Tric media. **A**. Bacterial optical density (OD_600_) was followed for 24 h in BHI (black line) and in SerFA-Tric (green line). In SerFA-Tric, growth resumed after a 10 h latency period. **B**. Fatty acid profiles. Newman strain in BHI (exponential culture, left), display endogenous fatty acids, and a SerFA-Tric culture at OD_600_ = 1 (15 h culture, right), comprising exclusively the 3 eFA. Arrow indicates endogenously produced *ai*15. eFA: 1, C14:0; 2, C16:0, 3, C18:1. Growth curves and profiles are representative of three experiments.

**Fig. S2.**
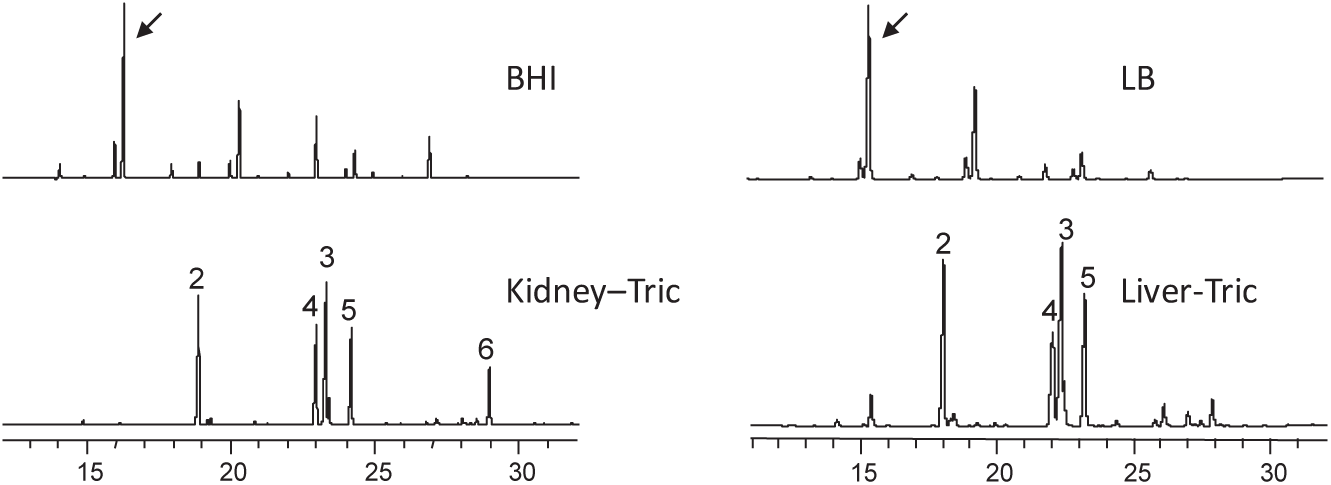
Fatty acid profiles of *S. aureus* grown in kidney or liver extract and triclosan. *S. aureus* USA300 strain was grown overnight in: BHI and LB, and BHI plus 3% pork kidney and LB plus 3% pork liver extracts to which Triclosan (0.5 µg/ml) was added. Organ preparation and extractions for gas chromatography are described (Materials and Methods; (1)). Organ fatty acids are: 2-C16:0; 3-C18:1; 4-C18:0; 5-C18:2; 6-C20:4. Results are representative of two independent experiments in each condition. *S. aureus* endogenous fatty acid *ai*15 is indicated by a black arrow.

**Fig. S3.**
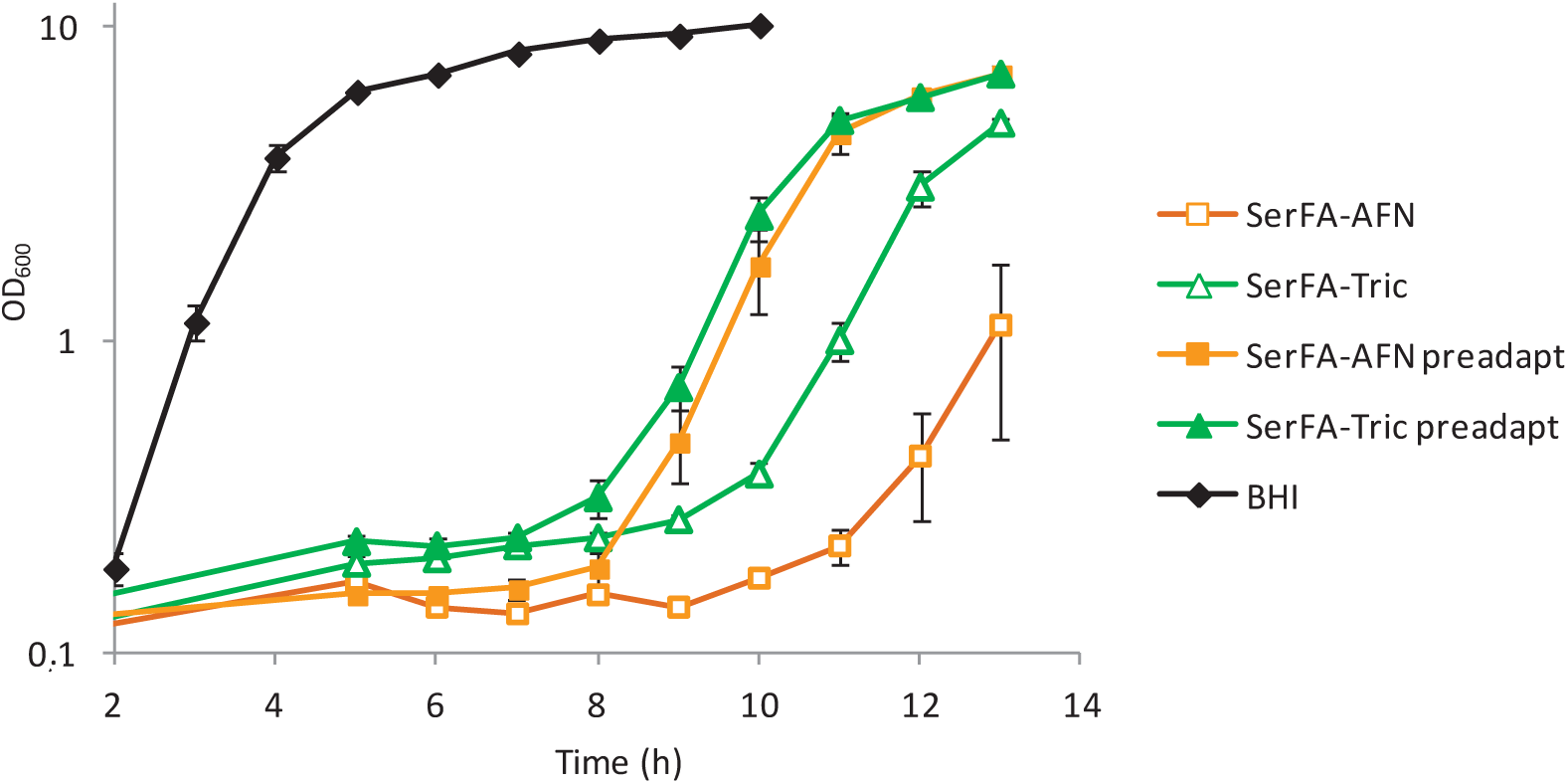
Impact of preadaptation in serum on *S. aureus* FASII antibiotic adaptation. Bacteria are generally exposed to the lipid-rich host environment prior to antibiotic treatment. Here, USA300 was precultured in BHI and in SerFA medium, and then challenged with either triclosan (green triangles) or AFN-1252 (orange squares). Growth of antibiotic-challenged cultures without preadaptation (empty markers) or with preadaptation (preadapt; filled markers) was followed by optical density (OD_600_) for 13 h. All FASII-antibiotic-treated cultures displayed exogenous fatty acid profiles at this time (available on reqques). Growth curves show the average and range of three biological replicates from one of two independent comparable experiments.

**Fig. S4.**
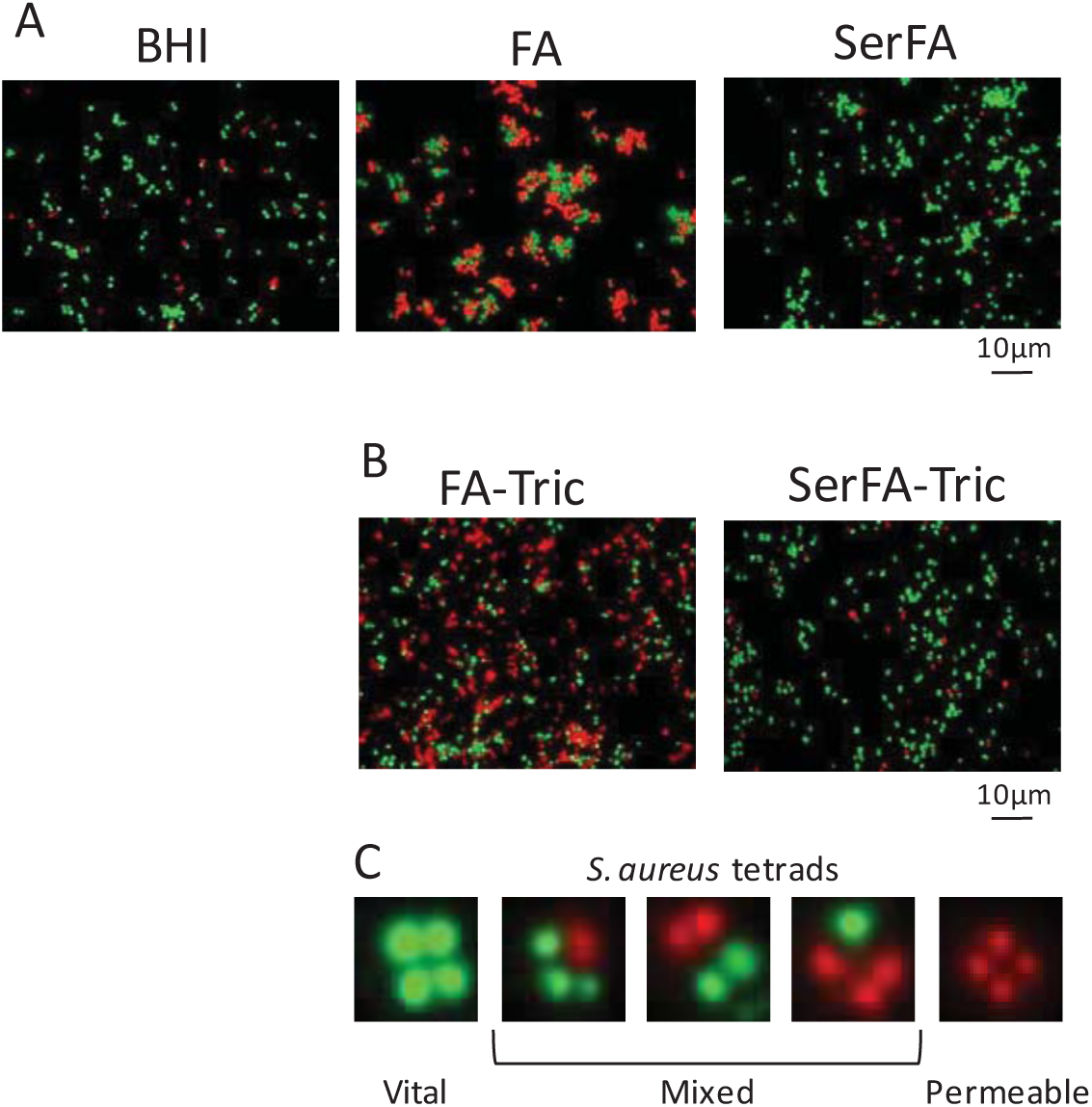
Non-aureus staphylococcal pathogens adapt to FASII antibiotic AFN-1252 by eFA incorporation. *S. aureus* USA300, *Staphylococcus epidermidis* ATCC 12228, *Staphylococcus haemolyticus* JCSC1435, and *Staphylococcus lugdunensis* N920143 were cultured in BHI and in SerFA-AFN medium. Fatty acid profiles from mid-log BHI cultures (left), and from 13-15 h SerFA-AFN cultures (right) are shown. Black arrow indicates endogenously produced *ai*15. eFA: 1, C14:0; 2, C16:0, 3, C18:1.

**Fig. S5.**
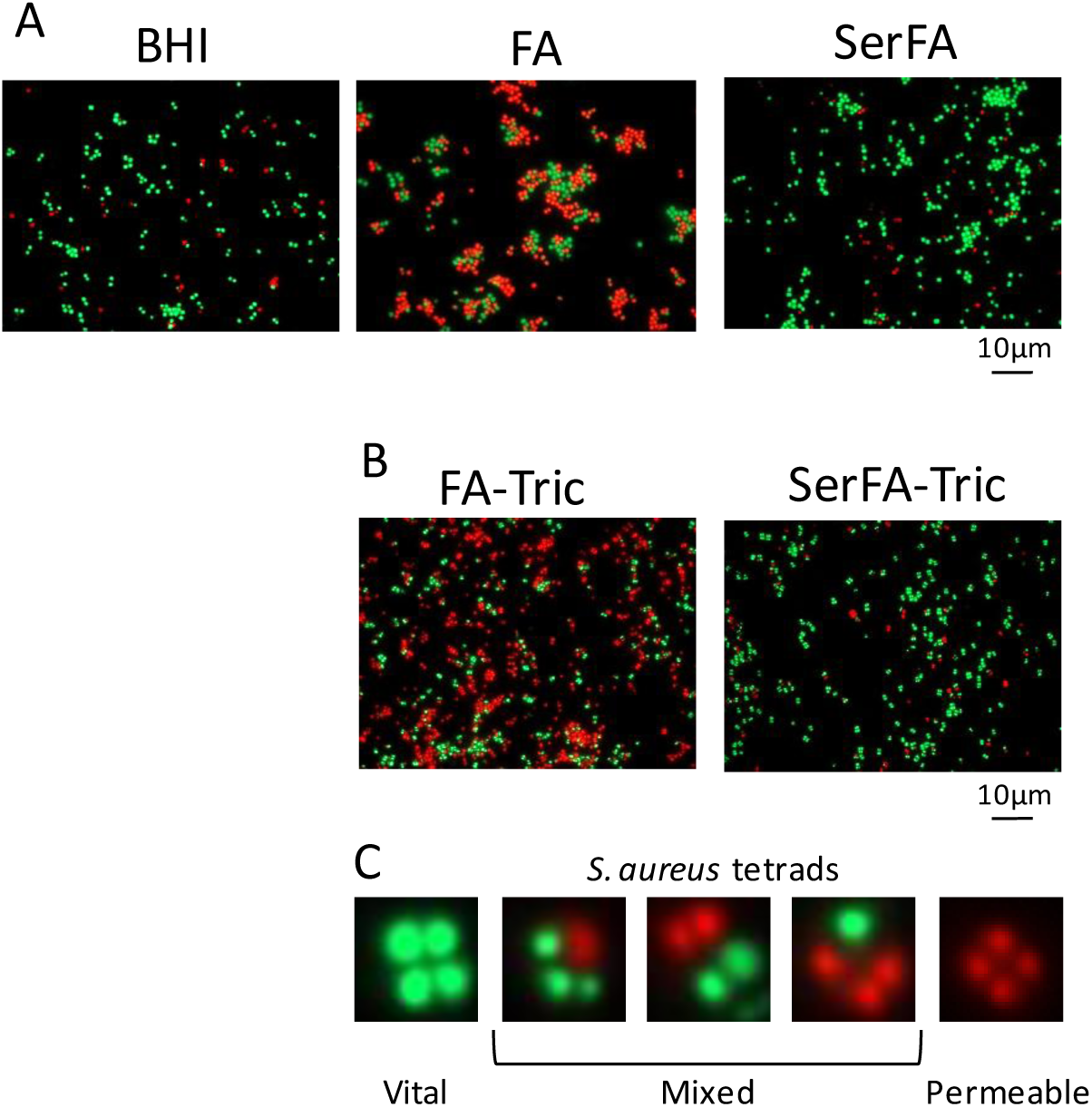
Serum improves *S. aureus* vitality during fatty acid and anti-FASII treatment. *S. aureus* USA300 was cultured in the following media, then stained with Syto9^TM^ (for cell vitality; green) and Propidium iodide (PI, for cell permeability; red), and visualized by fluorescence microscopy. **A,** BHI, FA, or SerFA media to OD_600_=0.5-1, and **B,** in FA-Tric or SerFA-Tric for 6 h. **C.** Tetrad clusters were frequent in antibiotic selections (shown for SerFA-Tric, and representing about 28% of total cells), and often comprised green and red cells (“Mixed”). Microscopy images were used to determine the proportions of vital, permeabilized and mixed-fluorescence cells (Fig. 2C).

**Fig. S6.**
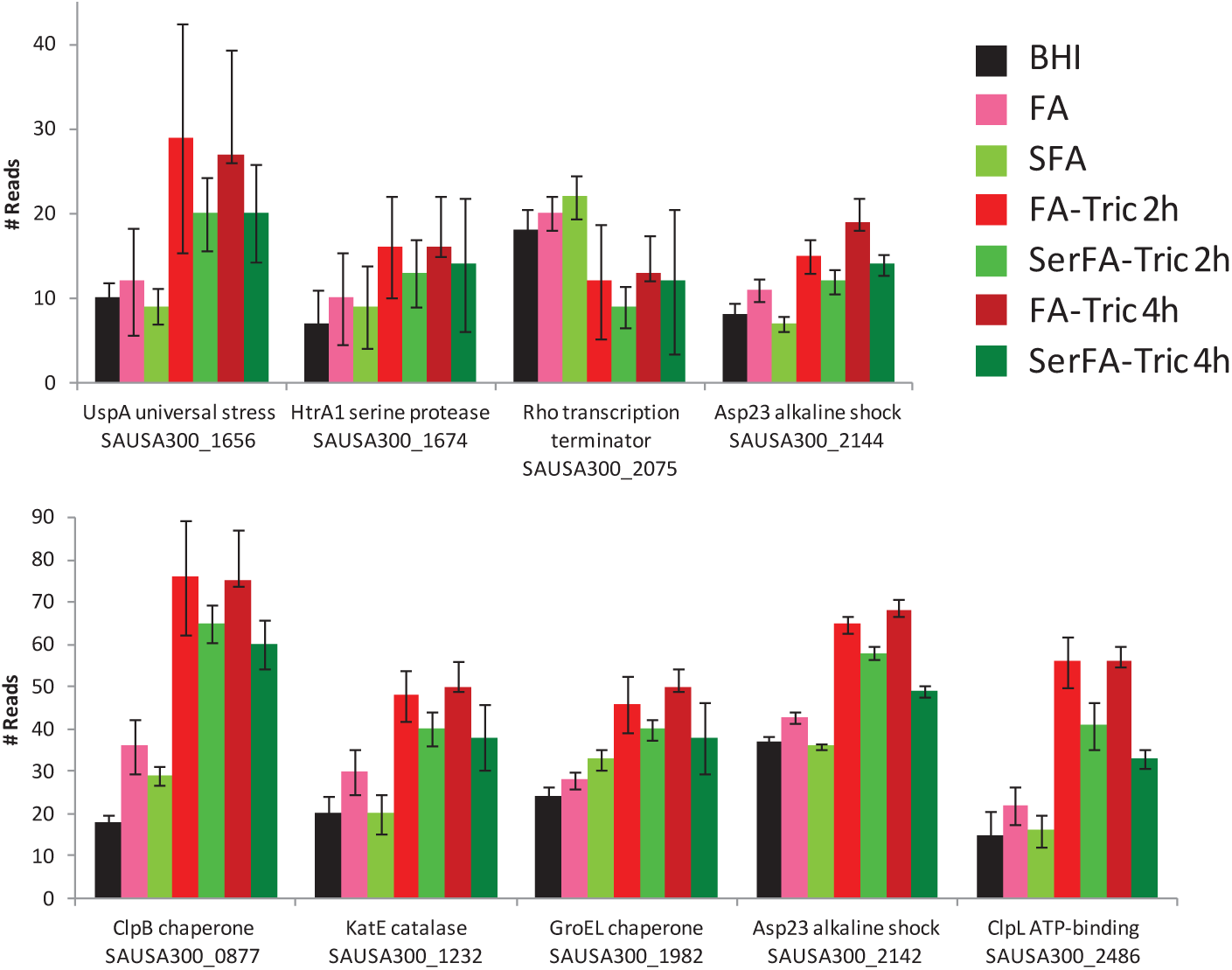
Proteomics of *S. aureus* responses to FASII antibiotic treatment. *S. aureus* USA300 *spa* was grown in BHI (black), FA (pink), SFA (light green) to OD_600_ = 1, in FA-Tric for 2 h (red) and 4 h (burgundy), and in SerFA-Tric for 2 h (mid-green) and 4 h (dark green) for proteomic studies (described in Materials and Methods). Proteins showing differences in bacterial stress response according to conditions are presented. The average number of reads and standard deviation are determined from quadruplicate independent samples. The full proteomic study is available at doi:10.17632/9292c75797.1.

**Fig. S7.**
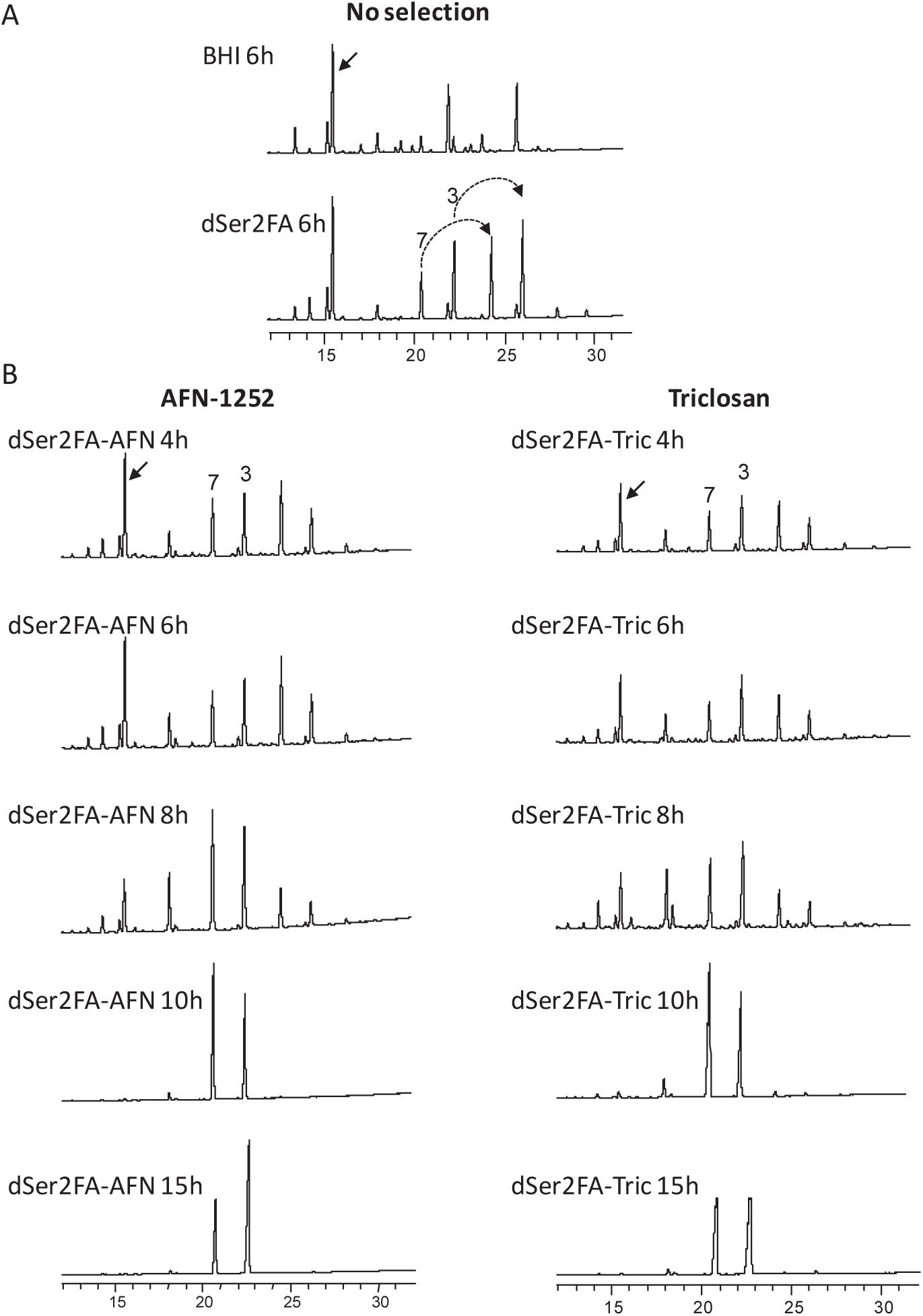
*S. aureus* USA300 incorporation of unsaturated fatty acids in non-selective and FASII-antibiotic-selective growth. Fatty acid profiles were determined by gas chromatography from cultures grown in dSer2FA (delipidated serum containing 17:1*tr* and 18:1*cis* (oleic acid)), neither of which are synthesized by *S. aureus*. AFN-1252 or triclosan was added to dSerFA preadapted cultures, and fatty acid profiles were analyzed at the indicated times. **A**, without antibiotic; **B**, with added AFN-1252 (left) and triclosan (right). The shift in fatty acid profiles in FASII-antibiotic-containing cultures is concomitant with growth restart at 8 h (Fig. S8). Black arrow, position of endogenously produced *ai*15. eFA used in experiment: 3, C18:1 *cis*; 7, C17:1*tr*. Dashed arrows, elongation (n+2) products of eFA.

**Fig. S8.**
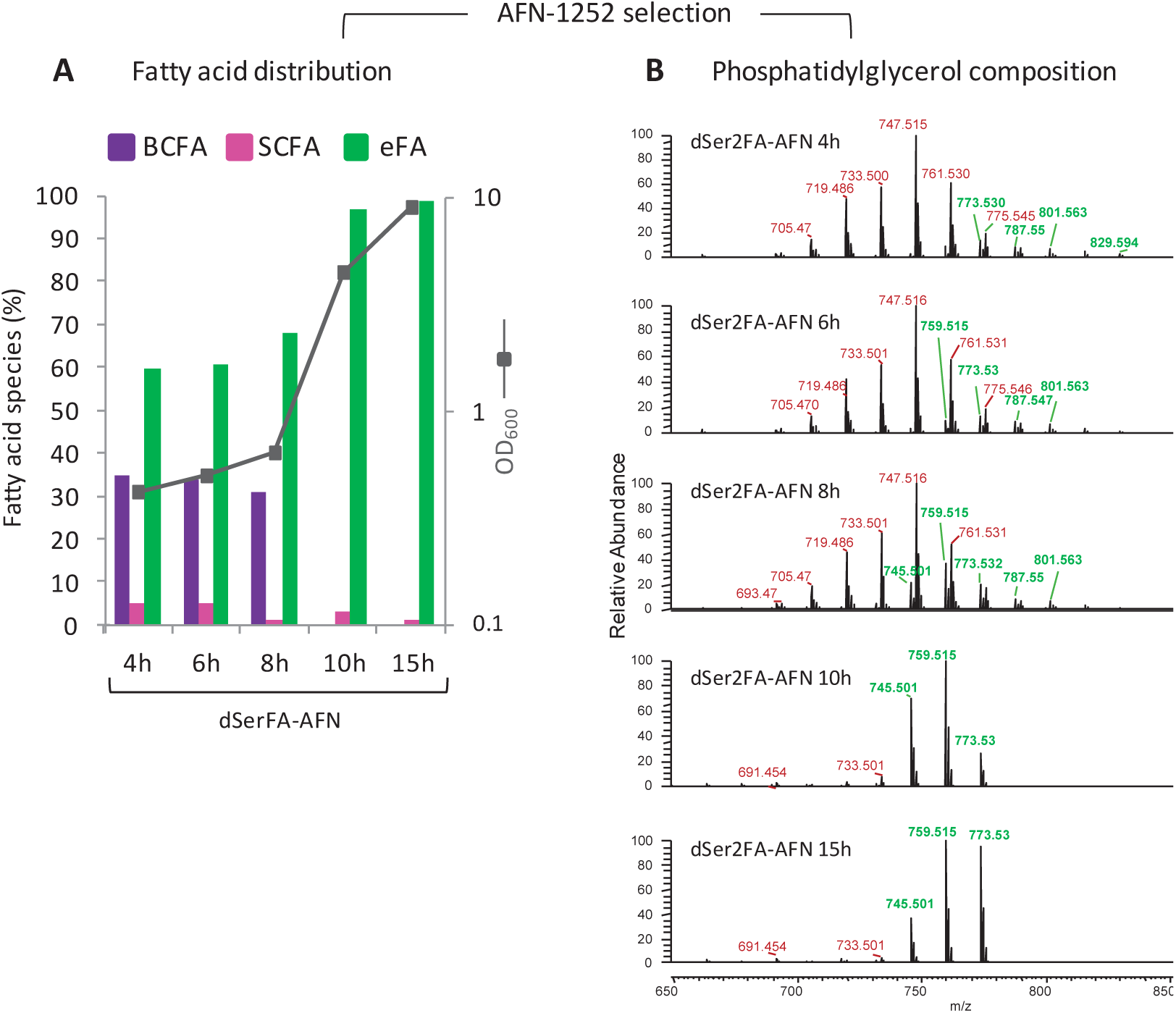
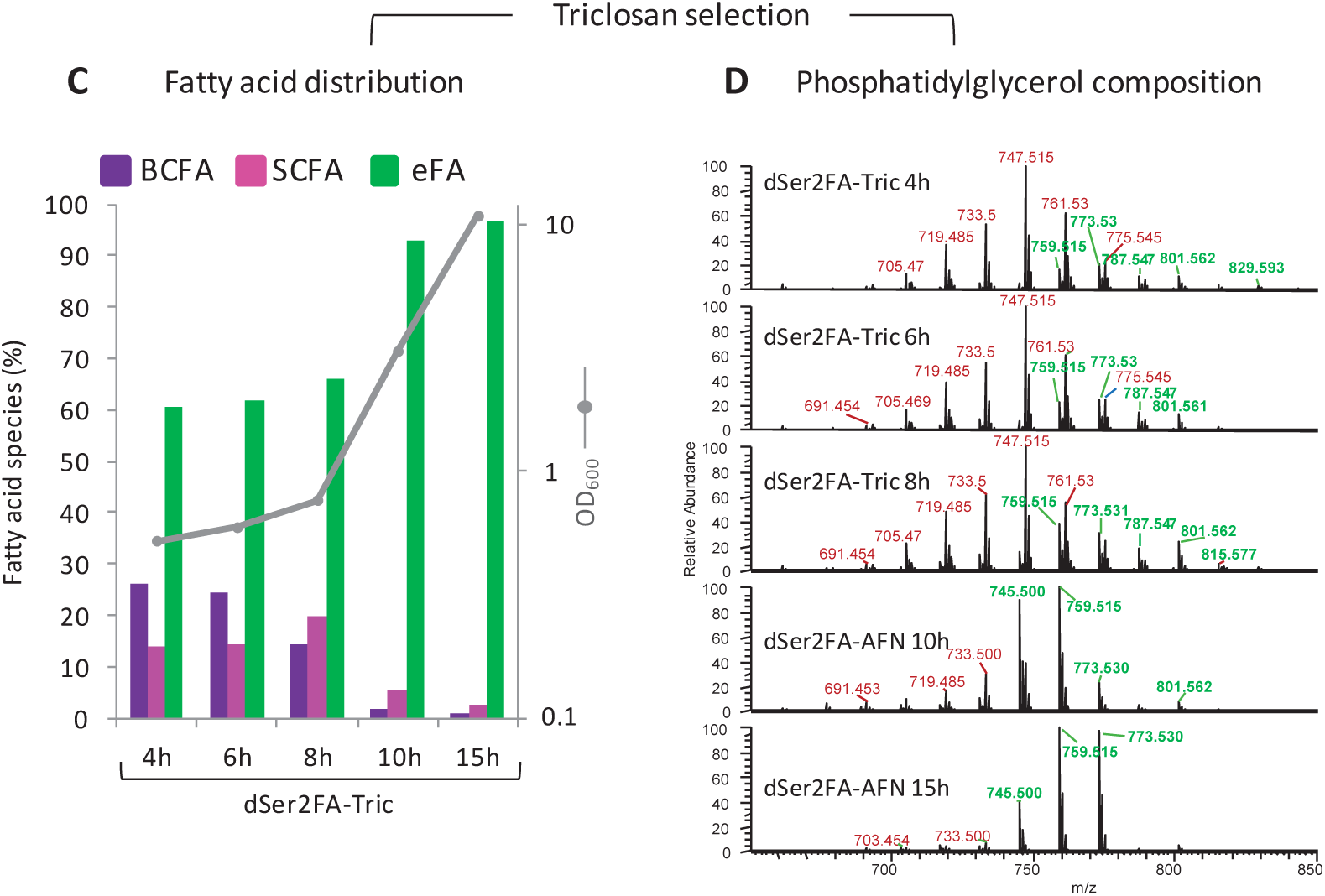
Kinetics of eFA incorporation in both phospholipid positions during FASII antibiotic selection. *S. aureus* USA300 dSer2FA pre-cultures and cultures were prepared with 10% delipidated serum (dSer) and C17:1*tr* and C18:1*cis.* AFN-1252, 0.5 µg/ml (**A** and **B**, preceding page) or Triclosan 0.5 µg/ml (**C** and **D**) was added at OD_600_ = 0.1 at the start of kinetics. **A** and **C.** Fatty acid composition at indicated time points is shown as the proportion of endogenous branched chain fatty acids (BCFAs C15, i15, *ai*15; purple), endogenous saturated (straight chain) fatty acids (SCFAs C18:0, C20:0; pink), and eFA (C17:1*tr* and C18:1*cis*, green); corresponding profiles are shown in Fig. S7. Grey curve represents OD_600_ readings at indicated times. **B** and **D.** Kinetics of phosphatidylglycerol (PGly) profile modifications during FASII bypass in FASII antibiotics. Masses are represented: red, PGly species with one eFA and one endogenous fatty acid; green, PGly species with eFA in both positions. Fatty acid composition of each major PGly mass is given in Table S3A (dSer2FA-AFN) and Table S3B (dSer2FA-Tric). BHI and dSerFA cultures (Fig. 4) were analyzed together with this experiment.

**Table S1.**
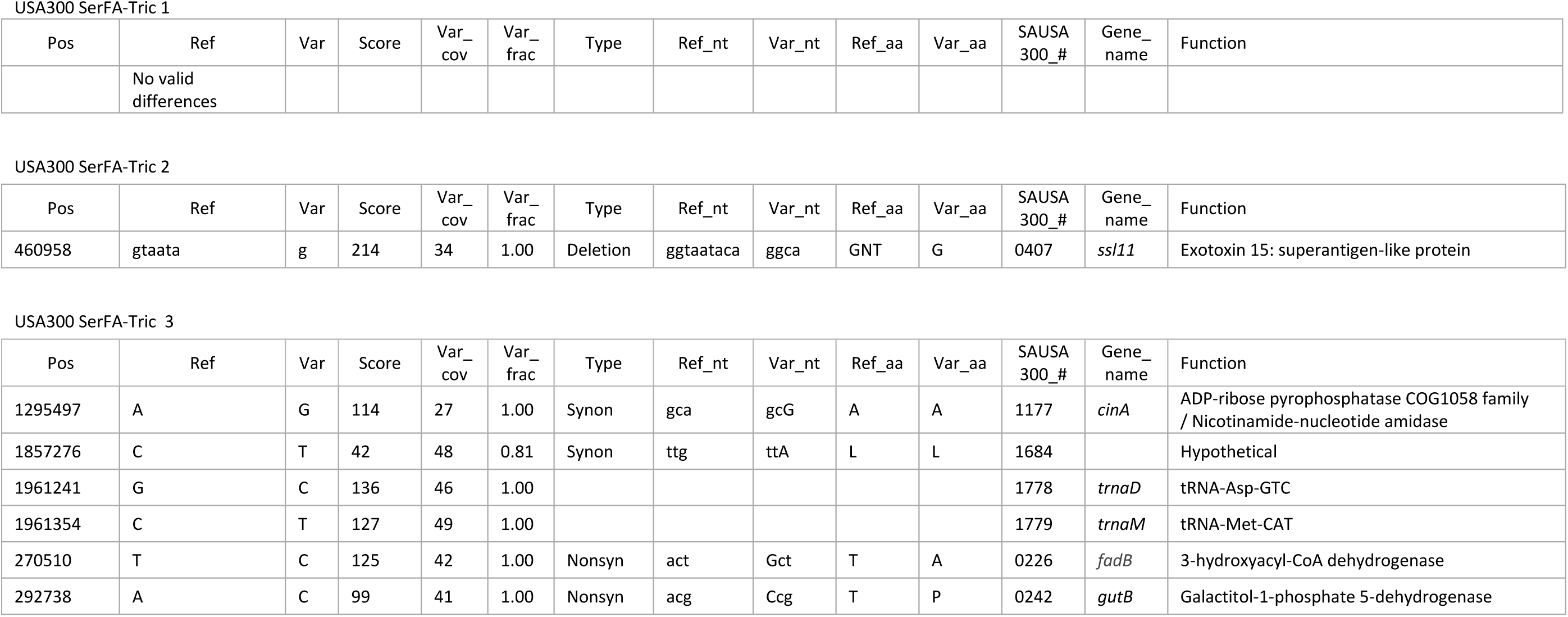
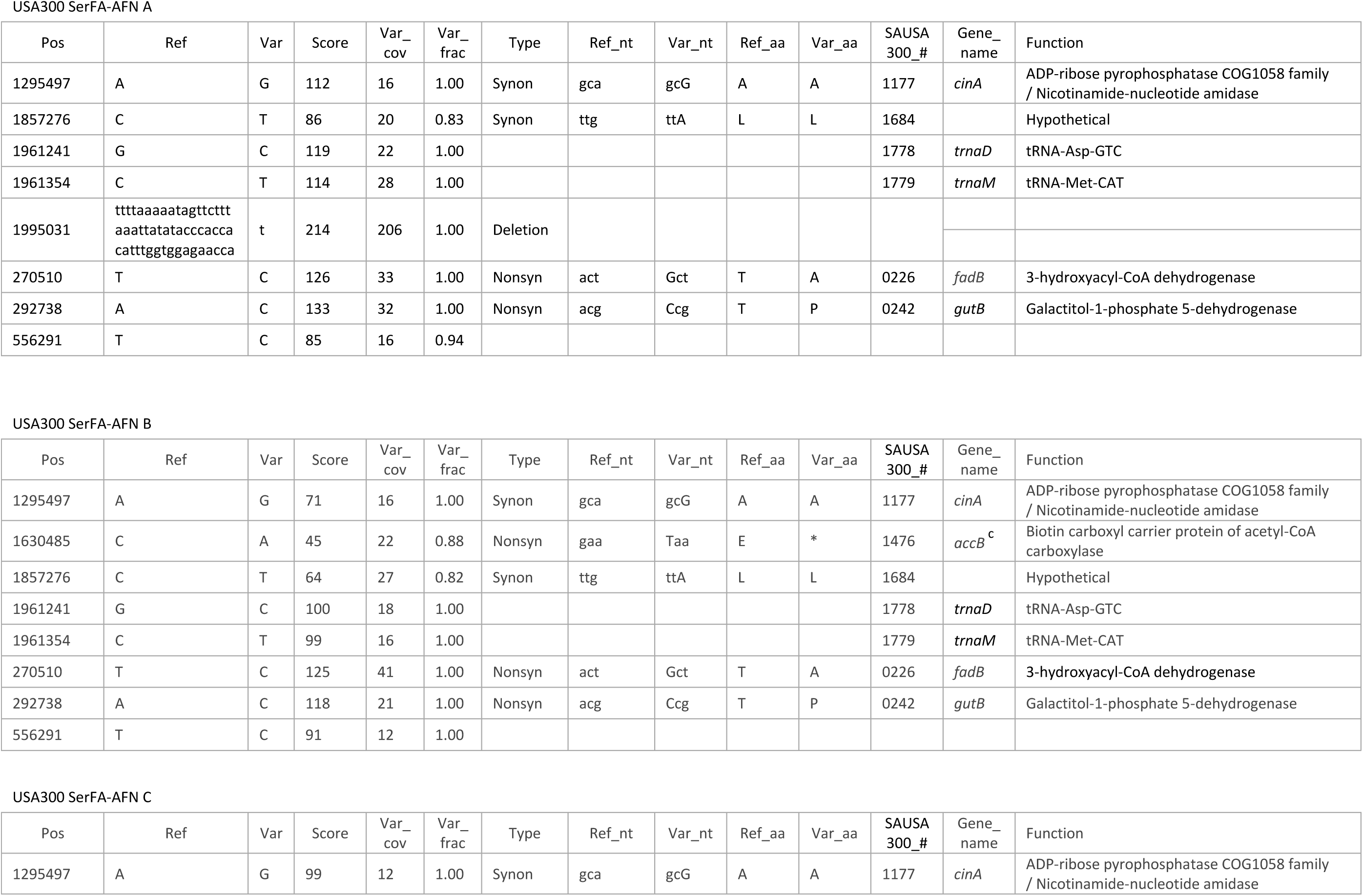
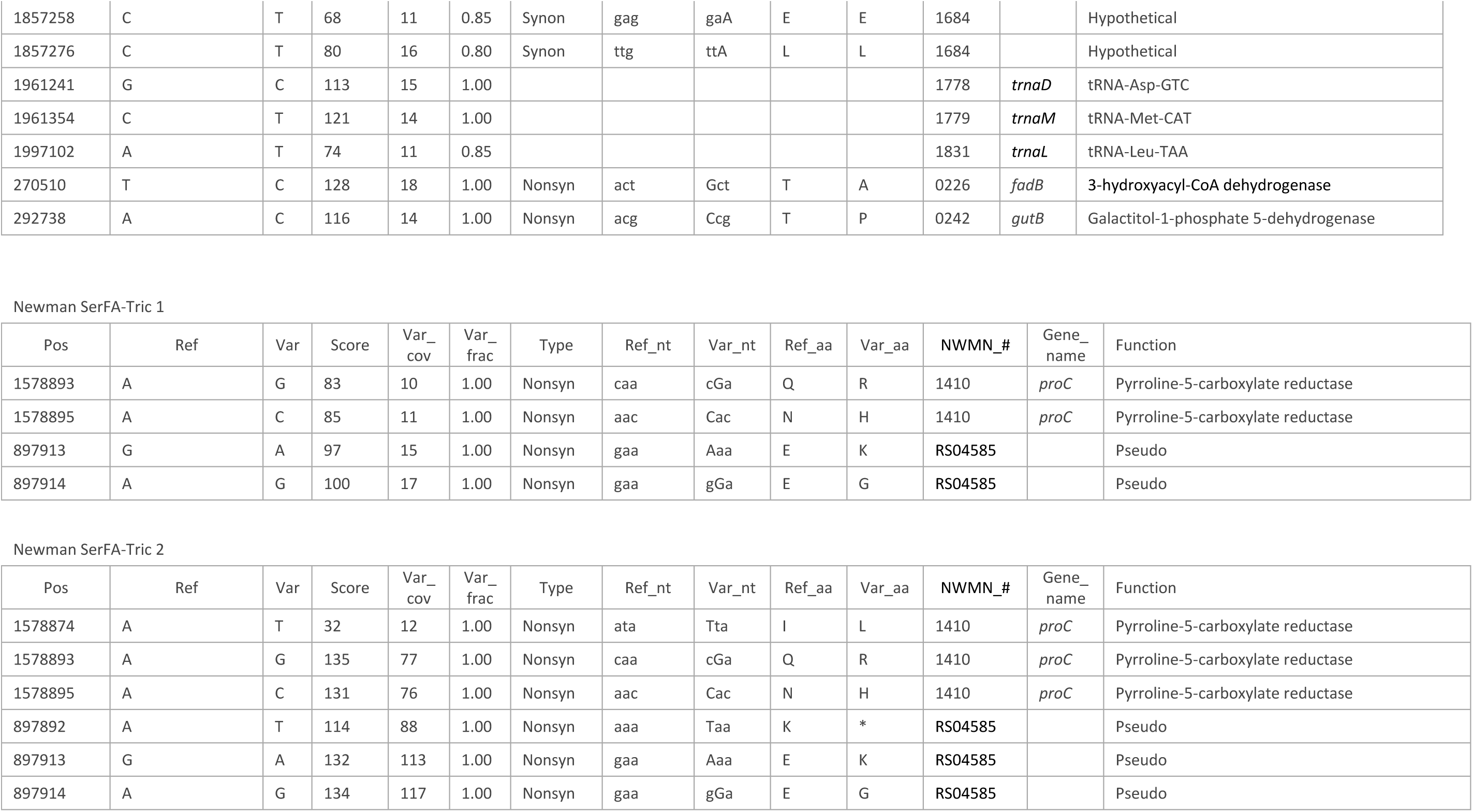
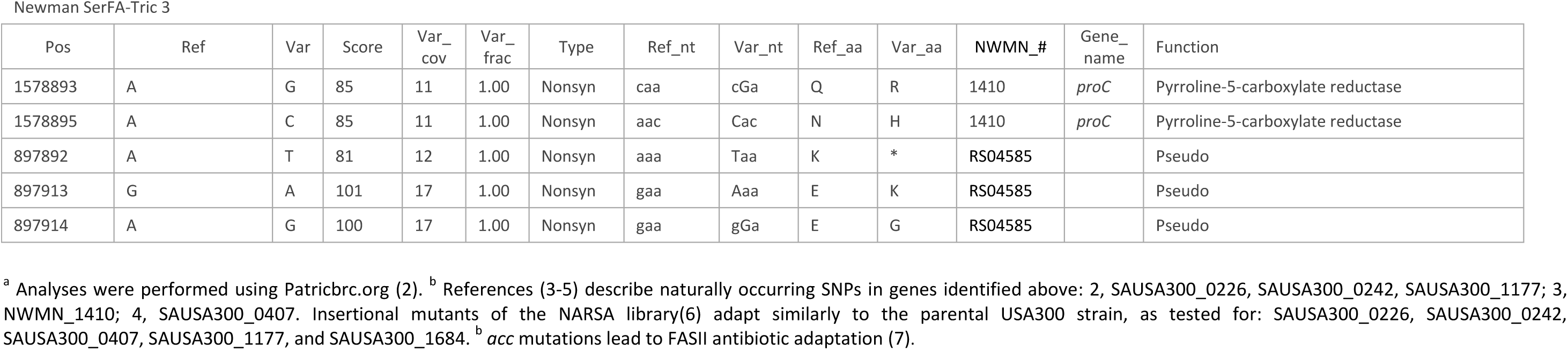
Sequence variations of SerFA-Tric and Ser-AFN *S. aureus* USA300 and Newman cultures.

**Table S2.** S. aureus USA300 fatty acid composition according to major PGly (phosphatidylglycerol) species in BHI and in medium containing dSer2FA (17:1tr and C18:1cis in delipidated serum). Lines shaded in green, fully exogenous PGly species; in pink, mixed endogenous/exogenous PGly species; in white fully endogenous PGly species. When 2 fatty acid constituents were identified for one mass, the minor species are in light print. In dSer2FA series: Note that all main PGly species comprise at least one eFA. See Fig. 4B for corresponding MS profiles.

| m/z | PGly | FA composition |
| --- | --- | --- |
| 679.453 | 29:0 | **14:0-15:0** |
| 693.469 | 31:0 | **15:0-15:0** |
| 707.483 | 31:0 | **16:0-15:0** |
| 721.500 | 32:0 | **17:0-15:0**<br>14:0-18:0 |
| 735.515 | 33:0 | **15:0-18:0** |
| 749.530 | 34:0 | **15:0-19:0** |
| 763.545 | 35:0 | **15:0-20:0** |
| 777.560 | 36:0 | **15:0-21:0** |
| 791.575 | 37:0 | 17:0-20:0<br>**15:0-22:0** |

**dSer2FA 6h culture**
| m/z | PGly | FA Composition |
| --- | --- | --- |
| 691.454 | 30:1 | **13:0-17:1** |
| 705.470 | 31:1 | **13:0-18:1**<br>14:0-17:1 |
| 719.485 | 32:1 | 14:0-18:1<br>**15:0-17:1** |
| 733.500 | 33:1 | **15:0-18:1** |
| 747.517 | 34:1 | **15:0-19:1** |
| 761.530 | 35:1 | **15:0-20:1** |
| 773.530 | 36:2 | **18:1-18:1** |
| 775.544 | 36:1 | **15:0-21:1**<br>18:0-18:1 |
| 787.547 | 37:2 | **17:1-20:1**<br>19:1-18:1 |
| 801.562 | 38:2 | **18:1-20:1** |
| 815.577 | 39:2 | **19:1-20:1** |
| 829.593 | 40:2 | **20:1-20:1** |

**Table S3A.** Fatty acid composition of PGly becomes fully exogenous during *S. aureus* USA300 treatment with anti-FASII AFN-1252 in medium containing dSer2FA-AFN (17:1*tr* and C18:1*cis* in delipidated serum).

|  |  |  | Time after AFN-1252 addition |  |  |  |  |
| --- | --- | --- | --- | --- | --- | --- | --- |
| m/z | PGly | Fatty acid composition | 4h | 6h | 8h | 10h | 15h |
| 691.454 | 30:1 | 13:0-17:1 | ++ | * | ++ | - | - |
|  |  | 12:0-18:1 | -- |  | ++ | ++ | ++ |
| 693.469 | 30:0 | 16:0-14:0 | ++ | ++ | ++ | * |  |
|  |  | 15:0-15:0 | ++ | ++ | + |  |  |
| 705.470 | 31:1 | 14:0-17:1 | ++ | ++ | ++ | ++ |  |
| 717.469 | 32:2 | 14:1-18:1 |  |  |  |  | ++ |
| 719.485 | 32:1 | 14:0-18:1 | ++ | ++ | ++ | ++ | ++ |
|  |  | 17:1-15:0 | ++ | ++ | ++ | ++ | + |
| 733.500 | 33:1 | 14:0-19:1 | ++ | ++ | ++ | - | -- |
|  |  | 15:0-18:1 | ++ | ++ | ++ | + | + |
|  |  | 16:0-17:1 | - | - | ++ | ++ | ++ |
| 745.500 | 34:2 | 17:1-17:1 | * | * | ++ | ++ | ++ |
| 747.517 | 34:1 | 15:0-19:1 | ++ | ++ | ++ |  |  |
| 759.515 | 35:2 | 17:1-18:1 | * | * | ++ | ++ | ++ |
| 761.530 | 35:1 | 15:0-20:1 | ++ | ++ | ++ |  |  |
| 773.531 | 36:2 | 17:1-19:1 | ++ | ++ | + |  |  |
|  |  | 18:1-18:1 | ++ | ++ | ++ | ++ | ++ |
| 775.544 | 36:1 | 15:0-21:1 | ++ | ++ |  |  |  |
|  |  | 18:0-18:1 | + | + |  | ++ | ++ |
| 787.547 | 37:2 | 17:1-20:1 | ++ | ++ | ++ |  |  |
|  |  | 19:1-18:1 | ++ | + | ++ |  |  |
| 789.561 | 37:1 | 15:0-22:1 | ++ | ++ | * |  |  |
|  |  | 17:0-20:1 | ++ | ++ |  |  |  |
| 801.562 | 38:2 | 18:1-20:1 | ++ | ++ | ++ |  |  |
| 815.577 | 39:2 | 19:1-20:1 | ++ | ++ | ++ |  |  |
| 817.558 | 39:1 | 18:1-21:0 | ++ | ++ | ++ |  |  |
| 829.593 | 40:2 | 20:1-20:1 | ++ | ++ | ++ |  |  |

**Table S3B.** Fatty acid composition of PGly becomes fully exogenous during S. aureus USA300 treatment with anti-FASII triclosan in medium dSer2FA-Tric (containing 17:1tr and C18:1cis in delipidated serum). Lines shaded in green, fully exogenous PGly species; in pink, mixed endogenous/exogenous PGly species; in white fully endogenous PGly species. All major PGly species (in bold) contain at least one eFA throughout. *, no available MS2 profile. Intensities were estimated from MS profiles as: ++ >50%; +, 20%<Intensity<50%; -, 5%<I<20%; --, I<5%. *, *, no MS2 available. When 2 fatty acid constituents were identified for one mass, the minor species are in light print. In red ‘++’, dominant PGly species (see Fig. S8B and S8D for corresponding MS profiles).

|  |  |  | Time after triclosan addition |  |  |  |  |
| --- | --- | --- | --- | --- | --- | --- | --- |
| m-z | PGly | Fatty acid composition | 4h | 6h | 8h | 10h | 15h |
| 691.454 | 30:1 | 13:0-17:1 | ++ | ++ | ++ | ++ | - |
|  |  | 12:0-18:1 | - | - | + | ++ | ++ |
| 693.469 | 30:0 | 16:0-14:0 | ++ | ++ | ++ |  |  |
|  |  | 15:0-15:0 | ++ | ++ | ++ |  |  |
|  |  | 13:0-17:0 | - | + | ++ |  |  |
| 703.454 | 31:2 | 13:1-18:1 |  |  |  | ++ | ++ |
| 705.470 | 31:1 | 13:0-18:1 | + | ++ | ++ | ++ |  |
|  |  | 14:0-17:1 | ++ | ++ | ++ | ++ |  |
| 717.469 | 32:2 | 14:1-18:1 |  |  | + |  | ++ |
|  |  | 17:1-15:1 |  |  | ++ |  | - |
| 719.485 | 32:1 | 14:0-18:1 | ++ | ++ | ++ | ++ | ++ |
|  |  | 17:1-15:0 | ++ | ++ | ++ | ++ | + |
| 731.484 | 33:2 | 15:1-18:1 |  | ++ | ++ | ++ | ++ |
|  |  | 17:1-16:1 |  | - | - | ++ | ++ |
| 733.500 | 33:1 | 14:0-19:1 | + | ++ | + | + | -- |
|  |  | 15:0-18:1 | ++ | ++ | ++ | ++ | ++ |
|  |  | 16:0-17:1 | - | + | + | ++ | ++ |
| 745.500 | 34:2 | 17:1-17:1 | * | ++ | ++ | ++ | ++ |
| 747.517 | 34:1 | 15:0-19:1 | ++ | ++ | ++ | + |  |
|  |  | 17:0-17:1 | -- | -- | - | ++ |  |
| 759.515 | 35:2 | 17:1-18:1 | ++ | ++ | ++ | ++ | ++ |
| 761.530 | 35:1 | 15:0-20:1 | ++ | ++ | ++ |  |  |
| 773.531 | 36:2 | 18:1-18:1 | ++ | ++ | ++ | ++ | ++ |
| 775.544 | 36:1 | 15:0-21:1 | ++ | ++ | ++ |  |  |
|  |  | 18:0-18:1 | ++ | ++ | ++ |  |  |
| 787.547 | 37:2 | 17:1-20:1 | ++ | ++ | ++ | ++ |  |
|  |  | 19:1-18:1 | ++ | ++ | + | ++ |  |
| 801.562 | 38:2 | 18:1-20:1 | ++ | ++ | ++ | ++ | ++ |
| 815.577 | 39:2 | 18:1-21:1 | + | ++ | ++ |  |  |
|  |  | 19:1-20:1 | ++ | ++ | ++ |  |  |
| 829.593 | 40:2 | 20:1-20:1 | ++ | ++ | ++ |  |  |

## Notes

http://www.ebi.ac.uk/ena/data/view/PRJEB24433

doi:10.17632/9292c75797.1

